# NETSCOPE: Information-Theory Based Network Discovery and Analysis

**DOI:** 10.1101/2022.04.18.488630

**Authors:** Tido Bergmans, Tousif Jamal, Aya Rezeika, Chih-Chia Hsing, Tansu Celikel

## Abstract

Biological systems are naturally described as networks, spanning molecular interactions, cellular circuits, and brain-wide functional connectivity. Despite the ubiquity of network data, workflows for inferring network structure and then applying comparable graph analyses across modalities remain fragmented. We present NETSCOPE, an open-source, multi-platform toolbox for information-theoretic network inference and analysis. NETSCOPE estimates pairwise statistical dependence with mutual information (MI), derives weighted adjacency matrices, removes likely spurious edges using shuffle-based thresholds, and prunes indirect connections using the data processing inequality (DPI). A key feature is the conversion of MI-based similarity into a metric space via (normalized) variation of information (VI), enabling weighted shortest-path and centrality analyses that require distance-like edge weights. We validate the toolbox on synthetic data with known ground-truth topology and by reconstructing published molecular networks in *Saccharomyces cerevisiae*. We further demonstrate cross-domain use cases in single-cell transcriptomic networks, cell-level anatomical maps, EEG connectivity, and resting-state fMRI. NETSCOPE runs in Python and MATLAB/Octave, and is compatible with Jupyter/Colab workflows.

## 1 Introduction

The brain’s capacity for information processing arises from interactions among networks at multiple scales - molecular and genetic regulatory networks, cellular circuits, and large-scale structural and functional connectomes. Although each of these network layers can now be measured with increasing precision, we still lack general-purpose tools for relating *how* higher-level network organization (structural and functional connectivity) emerges from coordinated interactions at lower levels (molecular and cellular). As a result, neuroscience and neurobiology often accumulate rich, modality-specific network datasets without a common analytical language for integrating them across scales.

Systems-theoretical analyses of molecular networks illustrate what becomes possible when biological mechanisms are treated explicitly as networks rather than as independent variables. Even as methods for reconstructing molecular interactions continue to mature, network-based approaches have already informed studies of genetic disorders [Lim et al., 2017], oncogenomics [Creixell et al., 2015], brain plasticity [Kole et al., 2017, 2018b,a] and cellular circuits [Huang et al., 2022, Zeldenrust et al., 2021, Huang et al., 2020]. Framing molecular mechanisms as networks shifts emphasis from differential expression of single genes to coordinated dependencies among sets of genes, helping reveal pathways, modules, and interactomes that better match how biological regulation is organized.

At the macroscale, structural and functional brain networks provide complementary views of neural organization. Structural connectivity, typically estimated from diffusion MRI (dMRI) and tractography, describes the physical substrate of long-range communication. Functional connectivity, estimated from fMRI, EEG, or MEG, captures statistical dependencies among signals that vary with context and cognitive demand. Network analyses of these modalities have exposed organizing principles such as modularity and hubs, clarified how distributed circuits support cognition and behavior, and highlighted connectivity disruptions associated with disorders including Alzheimer’s disease, schizophrenia, and epilepsy. Yet despite their shared “network” framing, molecular/cellular and brain-scale networks are often analyzed with different toolchains, different dependence measures, and incompatible representations, limiting cross-scale synthesis.

A key bottleneck is how edges are defined. Molecular network inference commonly starts by computing pairwise similarity between genes across samples [Mc Mahon et al., 2014, Jackson et al., 2020].and the chosen dependence measure strongly shapes the inferred topology. Pearson correlation is computationally convenient, but it is restricted to linear relationships and can be sensitive to outliers [Gibson et al., 2013]. More expressive alternatives, including deep learning approaches [Shu et al., 2021] and visualization/comparison toolboxes, such as GeNeCK [Zhang et al., 2019] can model complex structure, but they typically demand extensive training data, careful regularization to avoid overfitting, and substantial compute.

Mutual information (MI), rooted in information theory, offers an attractive complementary basis for edge definition. MI directly quantifies statistical dependence without assuming linearity or prescribing a parametric relationship between variables. In principle, MI is invariant to invertible re-parameterizations of each variable, making it appealing for cross-modality work where measurement scales and preprocessing differ. In practice, however, MI must be estimated from finite data, and estimator choice and preprocessing (e.g., discretization strategies) can influence numerical values, highlighting the need for transparent, standardized workflows that support robust estimation and validation rather than ad hoc, one-off implementations.

Across neuroscience, MI-based networks have been used to capture dependence structures that are poorly described by linear metrics. For example, in pediatric epilepsy fMRI, MI-defined functional edges improved prediction of IQ relative to correlation-based networks [Zhang et al., 2018]. In post-stroke depression, MI-based EEG networks revealed reduced inter-hemispheric connectivity that covaried with symptom severity [Sun et al., 2018]. MI has also been applied to characterize task-dependent network reconfiguration during working memory and decision-making [Liao et al., 2022] and it can better capture oscillatory synchronization and cross-frequency coupling in EEG/MEG than traditional linear measures [Vicente et al., 2011]. In molecular biology, MI-driven methods such as **ARACNE** [Margolin et al., 2006], **parmigene** [Sales and Romualdi, 2011], and **PANA** [Ponzoni et al., 2014] infer co-expression networks by estimating MI between gene pairs and applying strategies such as DPI-based pruning, parallel computation, or integration of prior knowledge. Despite their impact, these approaches often culminate in thresholded (binary) networks, which can obscure biologically meaningful variation in association strength and introduce sensitivity to arbitrary cutoffs.

This limitation is particularly consequential for downstream graph analysis. Many graph-theoretic measures are most interpretable, and in some cases only well-defined, when edge weights encode graded connection costs or distances (e.g., for shortest-path-based metrics such as average path length and betweenness centrality). Consequently, a practical MI workflow should support *weighted* networks and provide principled transformations from dependence strength (e.g., MI) to a distance-like quantity suitable for weighted-graph computations. One information-theoretic option is the Variation of Information (VI) [Shannon and Weaver, 1949], which can be used as a dissimilarity between variables; when paired with appropriate normalization or monotonic transformations, VI-based edge lengths preserve graded relationships while enabling weighted-network statistics without collapsing the network to binary structure.

Motivated by these needs, we introduce **NETSCOPE**, a plug-and-play Python/MATLAB toolbox for constructing, analyzing, and visualizing *weighted* biological networks from mutual-information-based dependence estimates. NETSCOPE provides end-to-end workflows for MI-based inference and feature extraction, supports pathway analysis and clustering, and exports results to common network-visualization software. The toolbox is openly available on GitHub (https://github.com/DepartmentofNeurophysiology/NETSCOPE) and is tested for use in Google Colab, Anaconda, MATLAB, and Octave, with optional MATLAB–Python interoperability via Oct2Py [Sales and Romualdi, 2011]. By standardizing MI-based network construction and enabling weighted graph analytics across modalities, NETSCOPE aims to make cross-scale and multimodal network analysis more reproducible, comparable, and clinically translatable.

## 2 Methods

NETSCOPE has a modular organization (Fig. 1). After data import, normalization, and entropy calculations, mutual information is calculated between any given node (e.g. transcript) in the network. “Network analysis” routines allow sparsification of the network by identifying indirect connections and provide tools for graph-theory-based network analysis. Users can optionally export the data for further analysis, e.g. in Gephi [Bastian et al., 2009], for custom visualizations.

**Figure 1:**
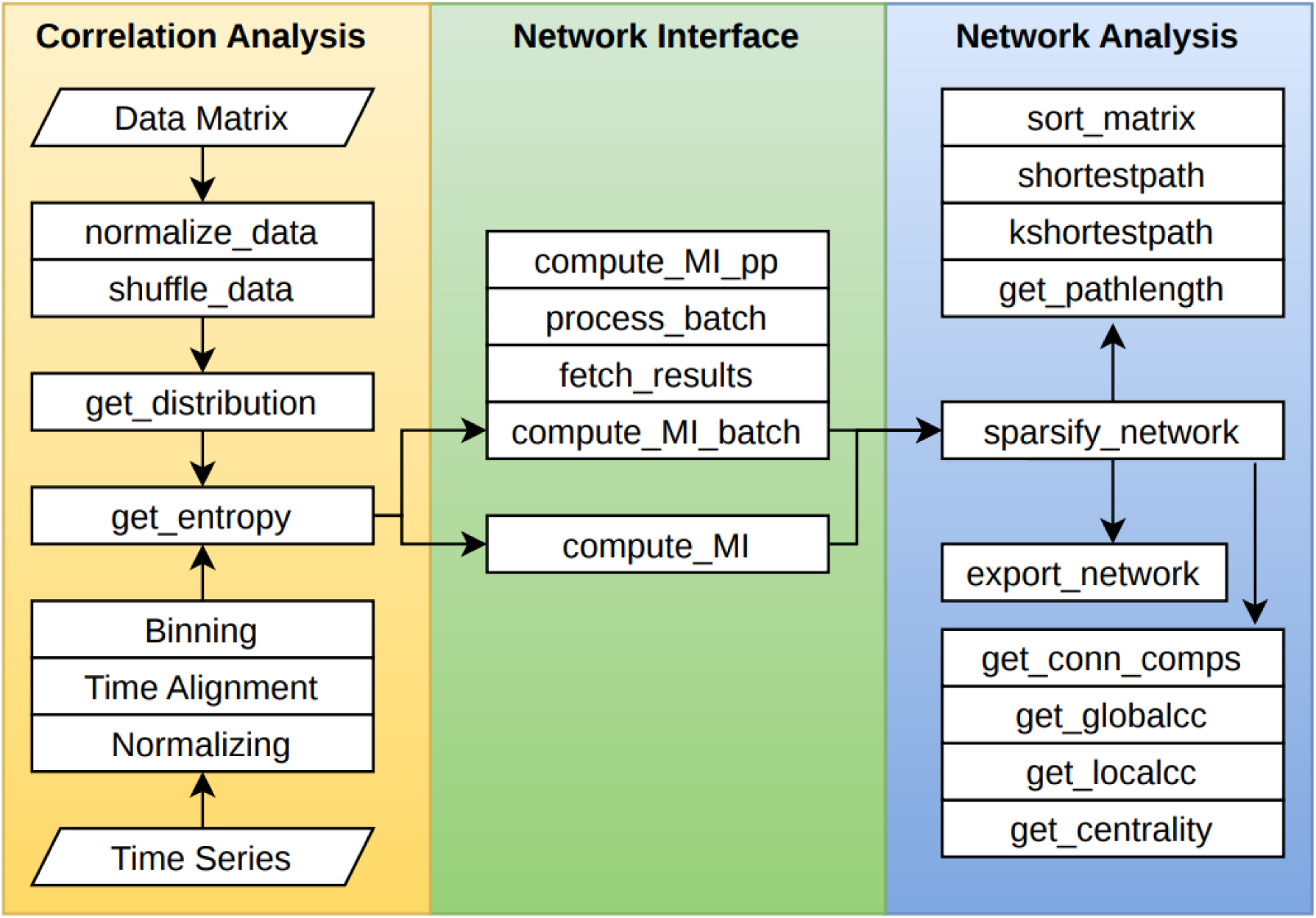
An overview of NETSCOPE functions and workflow. The data processing pipeline can be separated into three modules: data preparation, network inference, and network analysis. Function lists in each module and data processing hierarchy are shown.

The toolbox implements a data-processing pipeline that imports high-dimensional datasets organized as matrices, where columns represent samples (e.g., cells) and rows represent molecular targets (e.g., genes). From this input, NETSCOPE constructs weighted networks based on information-theoretic dependencies. Fig. 2 illustrates the workflow for transcriptional network construction. Beyond network inference, the toolbox provides functions to quantify net-work topology and identify hubs, clusters, and pathway structure. Although the terminology is transcription-focused, the framework is general and also applies to cellular networks in which nodes represent anatomical loci. A key methodological distinction of NETSCOPE from tools such as ARACNE and parmigene lies in its use of Variation of Information (VI) as a true distance metric. Although standard approaches rely on mutual information to infer pairwise associations, mutual information itself is not a metric and cannot directly support shortest-path computation. By transforming dependency structure into a metric space via VI, NETSCOPE enables weighted shortest-path analysis and principled evaluation of global communication pathways.

**Figure 2:**
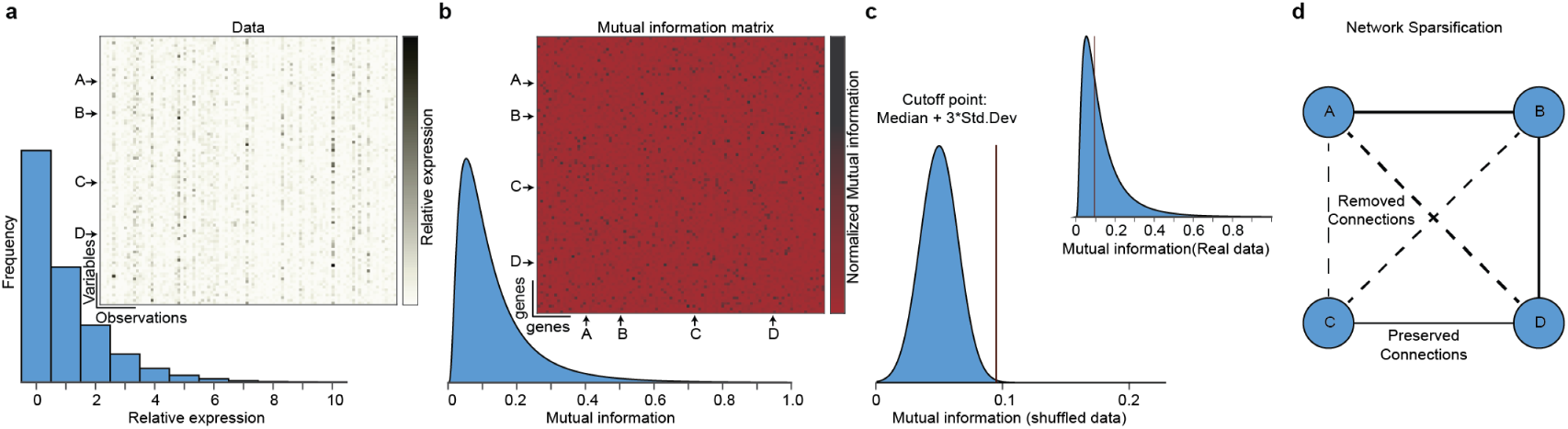
Basic processing steps for quantification of co-expression network from gene ex-pression data. **a** The pipeline starts with a data set organized as a matrix, e.g. RNA-sequencing data, where columns are samples (cells, repetitions) and rows are targets (e.g. transcript names). **b** From the expression data, the mutual information (MI) matrix is computed (see Materials and Methods for details). This matrix is interpreted as the adjacency matrix of a weighted, undirected network. **c** To establish a significance threshold for MI, a shuffle correction is performed for which MI matrices are computed from randomly shuffled gene expression data (see the section on “Shuffle correction”). A threshold is then based on the median and standard deviation of the shuffled MI distribution. **d** A network sparsification algorithm is applied to remove spurious correlations from the network. This algorithm iterates over the network connections, removing the edges that are the result of an indirect correlation (see the section on “Sparsification by data processing inequality”).

### 2.1 Network inference

The inference of gene co-expression networks is based on statistical dependencies between transcription levels of individual genes. Mutual information (MI) is used as a measure of correlation to describe these dependencies and quantifies the amount of information obtained about a stochastic variable through the information describing the other variable [Shannon and Weaver, 1949, Silvester]; see [Azarfar et al., 2018] for a recent expansion. It captures any type of statistical dependency, not just positive or negative linear relations, and is invariant under reparameterization. Mutual information between two genes is computed using the probability distributions of their respective transcription levels, as well as the joint probability distribution. The mutual information is then normalized by the joint entropy of the two genes. The joint entropy is a theoretical maximum for the mutual information so this constrains the normalized mutual information to the interval [0, 1]. The probability distributions of the genes are obtained by discretizing the transcription data using Sturges’ method [Sturges, 1926]. From these distributions, the entropy is calculated according to the equation:

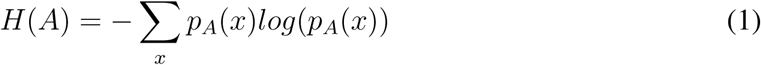

Where *H*(*A*) is the entropy of gene *A*, *p_A_* is the transcription distribution of *A*, and *x* is the discretized transcription level. From the same principle, the mutual information (MI) *I*(*A, B*) between genes *A* and *B* is calculated from the distributions *p_A_*, *p_B_*, and joint distribution *p_AB_*, and respective transcription levels *x, y*:

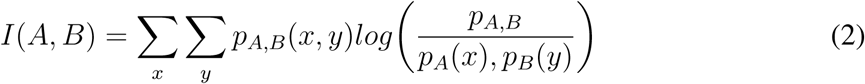

Computing pairwise MI between every set of two genes results in a matrix in which every entry (*A, B*) is the MI between genes *A* and *B*. This matrix is normalized with respect to the joint entropy of *A* and *B*, resulting in a symmetric network matrix with values between zero and one representing weighted connections in the gene co-expression network. The joint entropy can be calculated from the single gene entropies and MI:

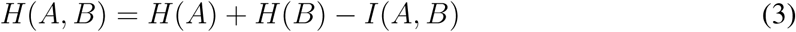

### 2.2 Shuffle Correction

In order to identify random correlations in the network, a significance threshold is established using shuffle correction. Shuffle correction is performed for each column across rows to preserve overall transcription per cell, but any correlations between genes are removed from the data. From this random gene expression matrix, the MI-based network is inferred according to the steps described in the previous section. This step is repeated *N* times (default: 10). With the distribution of MI values resembling a normal distribution, the significance threshold can be defined using the median and standard deviation of the distribution, e.g. *µ* + 3 ∗ *σ*, where *µ* is the median and *σ* is S.D. Connections with an MI value below this statistically defined cutoff point are considered random correlations and removed from the network.

### 2.3 Sparsification by Data Processing Inequality

The next step is to remove the connections from the network produced by indirect correlations. Indirect correlations appear when the transcription of two genes, e.g. *A* and *C*, is coregulated with a third gene *B* but otherwise independent. Since there is a correlation in transcriptional behavior between *A* and *B*, as well as between *B* and *C*, a spurious correlation will also be found between *A* and *C*, even though they do not interact and therefore should not be connected in the network. These indirect connections are removed using the principle of Sparsification by Data Processing Inequality [Margolin et al., 2006]. The Data Processing Inequality states that MI between *A* and *C* is limited by MI between *A* and *B*, MI between *B* and *C*:

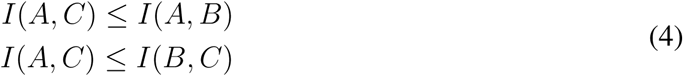

The sparsification is carried out by iterating over all first-order loops in the network (i.e. all trios of fully connected genes) and removing the weakest connection in the loop.

### 2.4 Network analysis

Various statistics, all of which are included in NETSCOPE, can be computed that help identify genes of interest and understand the overall structure of the network. At the node level, statistics such as degree, local clustering coefficient, and betweenness centrality can be used to identify genes that have an important role in the network, and potentially in transcriptional regulation. On a structural level, Dijkstra’s algorithm [Dijkstra, 1959] can be used to compute the (weighted) shortest path from one node to another, or the degree of separation (DOS), i.e. the minimum number of edges needed to get from one node to another. The degree of clustering in the network can be assessed by the global clustering coefficient, the small world coefficient, or PCA-related methods available in the toolbox. Finally, the toolbox contains functionality to export the network in GEXF format, a common data format used by network visualization software, such as Gephi [Bastian et al., 2009].

Since mutual information is a similarity (or “closeness”) metric, it cannot be used to describe distances between network nodes. To compute the shortest path from one node to another, the variation of information (VI) was used. The variation of information quantifies the information difference between two distributions [Meilă, 2007]; if MI describes how close two nodes are to each other in the network, then VI describes how far they are from each other. The variation of information can be calculated from the joint entropy and mutual information of a gene pair:

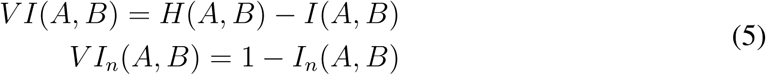

where *I_n_*(*A, B*) is the normalized mutual information. The maximum value for VI is 1 which is the case when MI equals zero and two genes are completely uncorrelated. Since two disconnected nodes in the network should have a “distance” of infinity (instead of 1), the distance metric in the network was defined as the ratio of VI to MI:

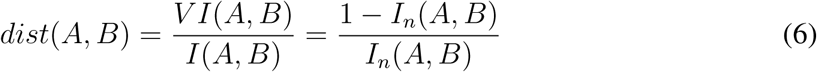

#### 2.4.1 Artificial EEG Network

In this study, we generate a synthetic EEG network as a benchmark to evaluate MI-based reconstruction. The ground truth network incorporates key characteristics observed in real EEG networks, including small-world properties, sparsity, and scale-free degree distributions. We then simulate EEG signals based on this network structure and reconstruct the connectivity using MI. By applying different thresholding strategies, we assess the effectiveness of MI in accurately recovering the ground truth network.

We generate a ground truth network consisting of 32 nodes, where each node represents an EEG channel. The network is defined as a binary adjacency matrix *G*, where an edge *G*(*i, j*) = 1 represents a functional connection between nodes *i* and *j*. The structure of *G* is designed to reflect key properties of real EEG networks:

- **Small-world properties**: The network maintains a balance between local clustering and long-range connections, ensuring efficient information transfer [Watts and Strogatz, 1998].
- **Sparsity**: EEG networks are typically sparse, with a connection density of around 15 − 20% of possible edges [Bassett and Bullmore, 2006].
- **Scale-free degree distribution**: Some nodes act as central hubs with significantly more connections than others, a feature commonly observed in functional networks [Barabási and Albert, 1999].

To prevent artificial alignment of the edges along the diagonal in visual representations, the node indices are randomly permuted while preserving the network structure. This ensures unbiased visualization of the connectivity matrix. To generate simulated EEG signals, we use a stochastic time series model that introduces dependencies between connected nodes. Each node represents a synthetic EEG signal, which is defined as:

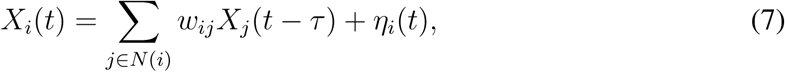

where:

- *X_i_*(*t*) is the signal at node *i* at time *t*,
- *w_ij_* is the interaction weight between connected nodes,
- *τ* is a small time delay to introduce causality,
- *η_i_*(*t*) is Gaussian noise.

We simulate each channel with randomly choosing weights(0-1) and a *τ* ∈ (0.5, 1.5*s*). Each channel generate a signal lasting for a total of 600 seconds. We inject a Gaussian noise ranging from 10% to 50%. The reconstructed MI-based connectivity was then compared with the known ground-truth network to quantify recovery performance.

### 2.5 EEG Data

All experiments have been collected after obtaining written consent from the participants according to the protocol approved by the Institutional Review Board at the Radboud University. Data were collected from 14 healthy right-handed participants aged 19 to 28 to investigate the effects of tactile and auditory stimuli on EEG recordings. The experiment utilized a passive tactile stimulus as the target and an auditory stimulus as a cue, with the latter presented prior to the tactile stimulus. Participants were equipped with EEG caps and electrodes, using electrolyte gel to maintain electrode impedance below 5 kΩ, and provided informed consent prior to participation. During the experiment, participants wore headphones, and a vibromotor was attached to the index finger of their dominant hand, while a fixation cross was displayed to maintain attention.

The auditory cue consisted of a 40 ms tone, followed by a tactile stimulus of 120 ms duration after a fixed interval, resulting in a total trial duration of 3500 ms. Each protocol consisted of 60 trials. Baseline recordings included auditory-only (AO) and tactile-only (TO) conditions, each with 60 trials per participant.

EEG data were acquired using two synchronized g.USBamp bioamplifiers (Guger Technologies, Graz, Austria) and recorded with BCI2000 (version 3.6) at a sampling rate of 512 Hz. Signals were filtered using a 2–100 Hz bandpass filter and a 50 Hz notch filter. The ground electrode was placed on the right earlobe and the reference on the left mastoid. A total of 32 Ag/AgCl electrodes were positioned according to the 10–10 system denoted as F_7_, FP_Z_, AF_8_, AF_3_, AF_4_, F_3_, F_z_, F_4_, FC_5_, FC_1_, FC_2_, FC_6_, T_7_, C_3_, C_Z_, C_4_, T_8_, TP_7_, CP_3_, CP_1_, CP_Z_, CP_2_, CP_4_, TP_8_, P_5_, P_1_, P_2_, P_6_, PO_Z_, O_1_, O_Z_, O_2_.

Preprocessing was performed using custom MATLAB scripts alongside EEGLAB and ERPLAB toolboxes. Data were re-referenced to the common average, and independent component analysis (ICA) was applied to remove artifacts. Components were classified using the ICLabel plugin [Pion-Tonachini et al., 2019], and artifactual components were rejected. The cleaned EEG signals were epoched from -1000 ms to 2000 ms relative to stimulus onset. Trials were grouped by stimulus type, and event-related potentials (ERPs) were computed for each condition.

Functional connectivity was estimated using the NETSCOPE toolbox by computing mutual information (MI) between all pairs of the 32 EEG channels. Four time windows were defined: (1) 250 ms before auditory stimulus onset (auditory baseline), (2) 250 ms before tactile stimulus onset (tactile baseline), (3) 250 ms after auditory stimulus onset (auditory evoked), and (4) 400 ms after tactile stimulus onset (tactile evoked). For each window, an MI matrix was computed. The top 10% of edges were retained to construct sparse functional networks, which were subsequently visualized using Gephi.

### 2.6 FMRI data

The dataset employed in this study consisted of preprocessed resting-state functional MRI (rs-fMRI) data from the Max Planck Institute Leipzig Mind-Brain-Body Dataset (LEMON, [Babayan et al., 2018]), including 136 healthy participants aged 20–30 years. The rs-fMRI data were parcellated into 183 regions using the Initial Parcellation Atlas (iPA), derived via voxel-level clustering constrained by macro-regions such as the frontal, parietal, occipital, and temporal lobes, as well as the insula and subcortical structures.

The NETSCOPE toolbox was extended to incorporate confound regression, enabling the extraction of clean timeseries by removing motion-related and physiological noise using a general linear model. Mutual information (MI) connectivity matrices were computed for each participant based on these cleaned timeseries, capturing nonlinear dependencies between all region pairs.

Individual MI matrices were averaged across participants to obtain a group-level MI connectivity matrix. This matrix was compared to group-averaged functional connectivity (FC) matrices based on Pearson’s correlation as provided by [Jimenez-Marin et al., 2024], allowing for a direct comparison between linear and nonlinear measures of functional connectivity.

To systematically compare connectivity patterns, an edge overlap analysis was performed. The top 10% of connections from the Pearson’s correlation-based FC matrix were selected to define significant edges. Both FC and MI matrices were then binarized, assigning a value of 1 to FC edges and 2 to MI edges. The matrices were summed to generate an edge overlap matrix, where values of 1, 2, and 3 indicated FC-only, MI-only, and overlapping edges, respectively.

To further evaluate the robustness of the NETSCOPE framework, a receiver operating characteristic (ROC) analysis was conducted using multiple thresholds (1%, 5%, 10%, 20%, 25%, and 50%) applied to the FC matrix. Additionally, Degree centrality, and node distance dendogram was computed for both weighted and unweighted networks to assess how edge weighting influences network topology.

## 3 Results

In the following two sections, we assess the quality of networks that were constructed using the toolbox. First, we show the performance of NETSCOPE with various discrete datasets. The network was constructed from artificial (synthetic) data with a known ground-truth network organization. Expression profiling was performed on the synthetic data, and the underlying network was inferred using NETSCOPE. Then we applied the NETSCOPE to construct network from RNA-sequencing data from Saccharomyces cerevisiae, a species of yeast whose genome has been studied extensively [Münzner et al., 2019, Dai et al., 2018, Janjić et al., 2014]. The resulting network was compared to five previously found co-expression networks in S. cerevisiae and a network pathway analysis was performed to identify potential genes of interest. Secondly, we show that we can use NETSCOPE for creating networks using continuous time series data. We created an artificial EEG like data with 32 channels and evaluating the performance against the ground truth data. Then we apply it to previously published data LEMON, and EEG data collected from experiment.

### 3.1 Ground truth network with discrete data

A topologically simple, undirected, unweighted ground truth network was created and used to generate synthetic expression data. The ground truth network consists of 100 disjoint graphs of 10 nodes connected in a linear pattern. From a random seed, expression data were generated that preserve this topology. The data was fed into the data processing pipeline to infer a co-expression network. The resulting network was shuffle corrected with a cut-off point of *µ*+3∗*σ*, sparsified and then compared to the ground truth network to assess its accuracy and to evaluate the limits of the method in terms of minimum data dimensionality. This was repeated by varying the number of samples in the expression data and the levels of noise added to the expression data. The number of samples ranged from 100 to 8000 and the noise was sampled from a normal distribution with a standard deviation of 10% to 50% of the mean. To assess the quality of the inferred networks as a function of data density and the number of samples, the true/false positive rates were used, where a false positive is a connection found between two nodes that were not connected in the original network. The results show that the true positive rate (TPR) increased with the number of samples used for network reconstruction. Notably, NETSCOPE maintained robust performance even under substantial noise. At a noise level of 50%, the method achieved a TPR of approximately 84% with as few as 100 samples, demonstrating resilience to high measurement variability.

Optimal reconstruction accuracy was achieved at lower noise levels (10%) in combination with larger sample sizes (≥ 1000), where the true positive rate approached its upper bound. These results demonstrate that increasing sample size substantially improves recovery of the underlying network topology, while the MI-based inference framework retains strong performance even under elevated noise conditions.

As expected, overall network quality scaled positively with the number of samples and declined with increasing noise amplitude (Fig. 3a), reflecting the balance between data density and signal degradation in network reconstruction. Distributions of MI values in the 8000 sample networks as a function of the degree of separation (DOS) in the original network revealed that the directly connected nodes have a median MI *>* 0.2 for all noise levels, with MI decaying as the distance between nodes increases (Fig. 3b). The decay rate increases with the amount of added noise, and disconnected nodes always have **MI** around zero.

**Figure 3:**
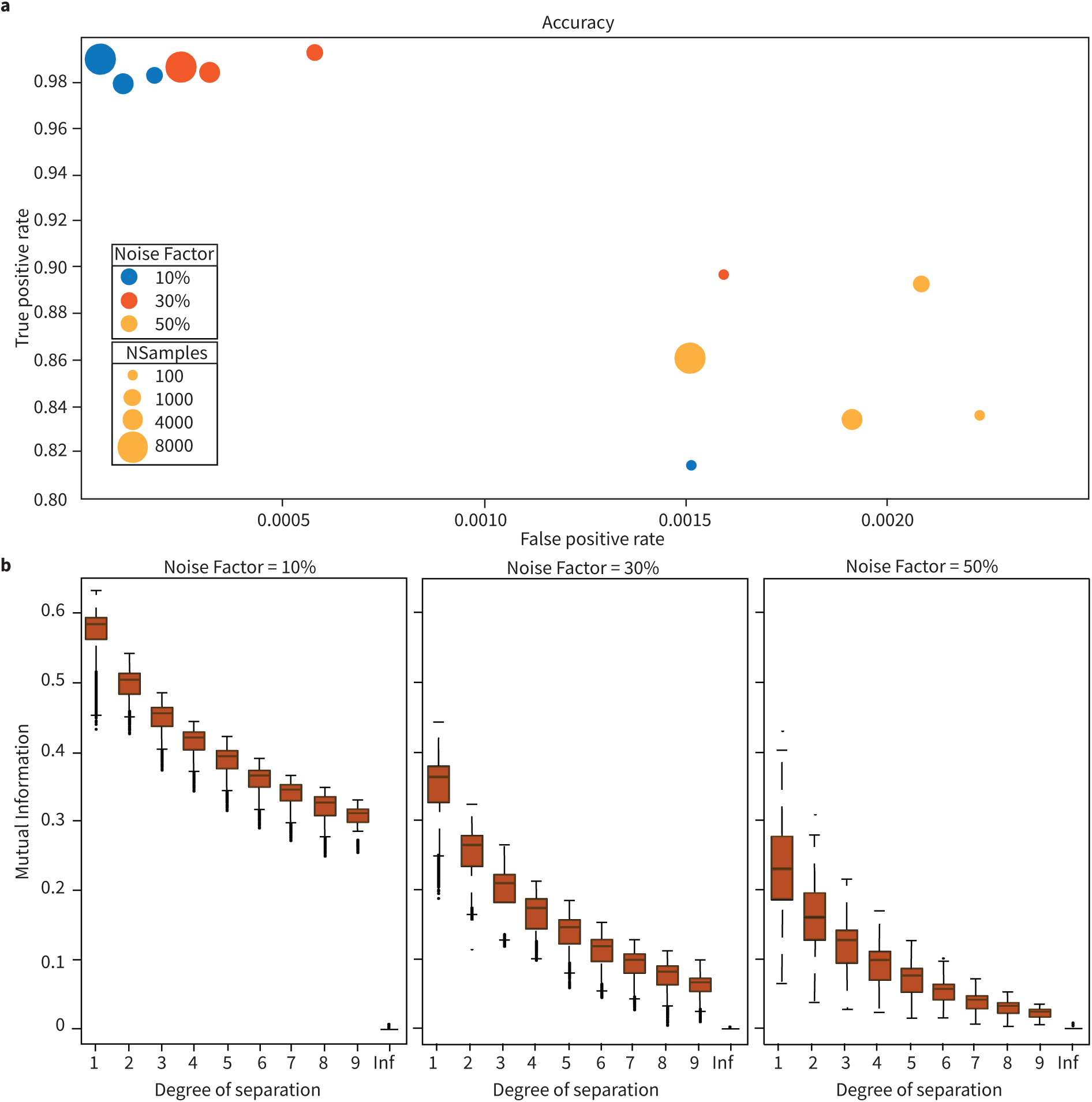
Networks constructed from synthetic gene expression data. Networks constructed from synthetic gene expression data. Networks were constructed from synthetic expression data with varying numbers of samples and variable amounts of noise added. **(a)** Accuracy of constructed networks when compared to their ground truth respective. Marker color denotes noise levels (red=10%, blue=30%, yellow=50%) and marker size denotes the number of samples used (100, 1000, 4000, 8000). **(b)** MI as a function of separation (DOS) in the networks. DOS is the number of edges between a pair of genes; a value of 1 means they are directly connected; a value of ‘inf’ means there is no possible path between them.

### 3.2 S. cerevisiae networks

To test the performance of NETSCOPE with biological systems, networks were constructed from RNA-sequencing data from S. cerevisiae [Ziemann et al., 2019]. A co-regulatory network was constructed and filtered after shuffle correction from 7126 genes sequenced in 672 samples. The subsets of this network were compared to five corresponding co-expression networks found in previous studies, which are listed in Table 1. The network data were made available by YeastNet *v*_3_ [Kim et al., 2014].

**Table 1:** Sources of the co-expression networks in Saccharomyces cerevisiae.

| No. | Reference | Title |
| --- | --- | --- |
| 1 | Mitchell et al. 2009 [Mitchell et al., 2009] | Adaptive prediction of environmental changes by microorganisms. |
| 2 | Wade et al. 2009 [Wade et al., 2009] | The Snf1 kinase and proteasome-associated Rad23 regulate UV-responsive gene expression. |
| 3 | Shalem et al. 2011 [Shalem et al., 2011] | Transcriptome kinetics is governed by a genome-wide coupling of mRNA production and degradation: a role for RNA Pol II. |
| 4 | Yona et al. 2012 [Yona et al., 2012] | Chromosomal duplication is a transient evolutionary solution to stress. |
| 5 | Orlando et al. 2008 [Orlando et al., 2008] | Global control of cell-cycle transcription by coupled CDK and network oscillators. |

As expected, compared to the synthetic data, the MI values were significantly lower in the yeast networks and the median value of normalized MI in each of the 5 networks was less than 0.05 (Fig. 4a). To numerically compare NETSCOPE’s network reconstruction against previously published network structure in yeast, we calculated the receiver operating characteristics (ROC). Because ROC is sensitive to noise level, we calculated three thresholds to binarize the MI matrix (Fig. 4a; three candidate thresholds established by shuffle correction are shown in color) and quantified the successful identification (i.e. true positive) against the false positive ratio (Fig. 4b). The expected results showed that with the threshold inversely proportional to the true and false positive discovery rate; with the increased threshold both discovery rates are reduced. Conservatively ∼ 60% of all previously identified networks could be correctly discovered using NETSCOPE with a false discovery rate of ∼ 20% (Fig. 4) using the conservative normalized MI threshold (i.e. 11 ∗ *σ*). The discovery rate improves to *>* 80% with lower thresholds (e.g. 7 ∗ *σ*) at the expense of an increased false discovery rate.

**Figure 4:**
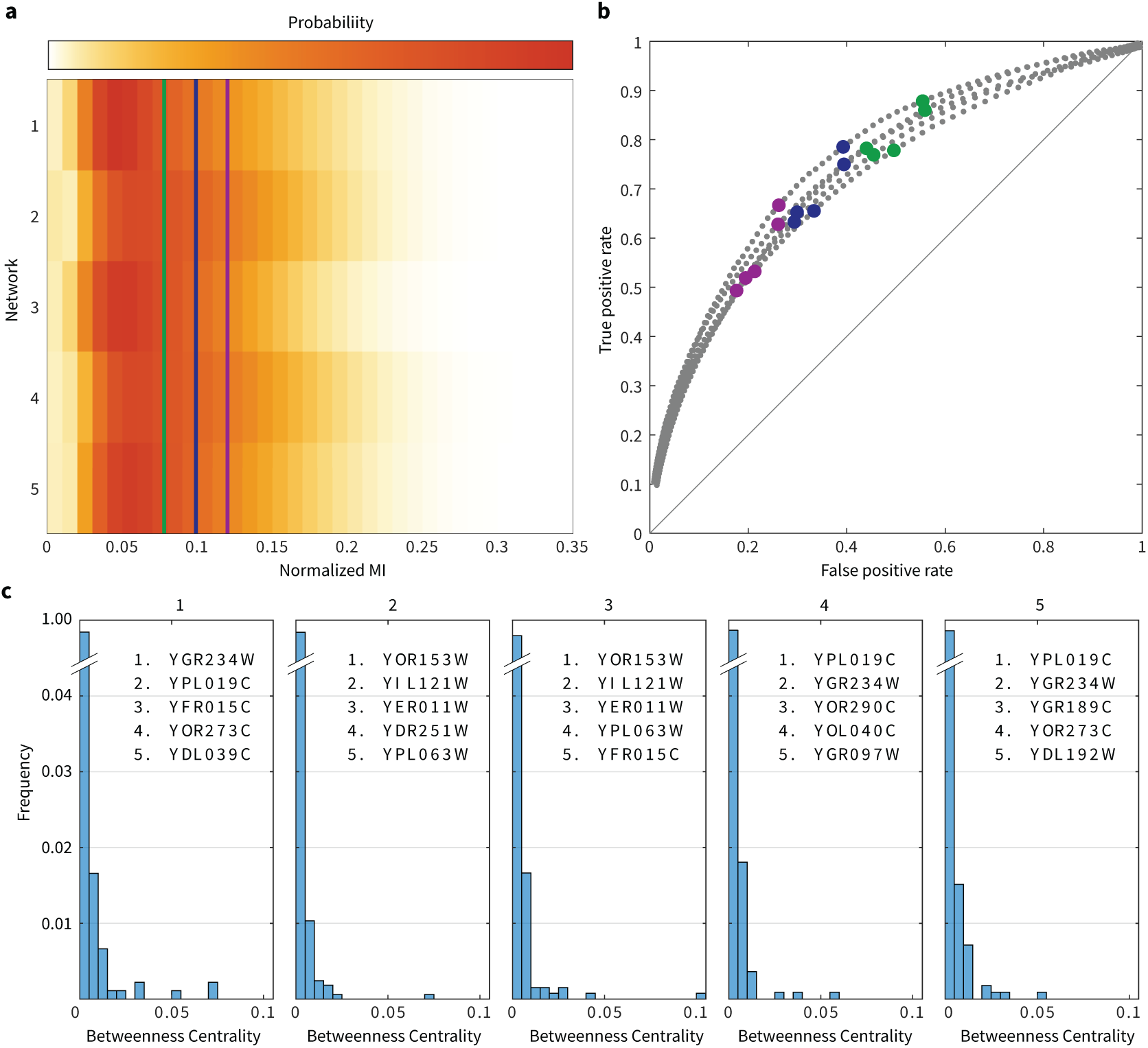
Five gene co-expression networks in S. cerevisiae constructed from RNA-seq data. **(a)** Distributions of MI values of the constructed networks. The three candidates’ *µ* plus 7 (green), 9 (blue) and 11 (purple) times *σ* of the shuffled MI distribution are shown. **(b)** Receiver operating characteristic (ROC) curve. True and false positive rates show the similarity between the constructed networks and previously established networks (see Table 1) for different significance thresholds. The colored data points correspond with the thresholds shown in **a**. **(c)** Distributions of the nodes’ betweenness centrality in the five networks. Every plot lists the top five genes with the highest betweenness centrality. Genes that are preserved across different networks are bold-printed.

NETSCOPE differs from previous approaches to MI-based molecular network inference (ARACNE [Margolin et al., 2006], parmigene [Sales and Romualdi, 2011] and PANA [Sales and Romualdi, 2011]) as such it allows weighted network analysis for pathway discovery (see the variation of information in Methods). The direct utility of identification of weights in the network is that it allows computation of the shortest path among any given two targets within the network, hence expanding the utility of MI analysis to pathway discovery. This approach can be expanded to the analysis of each target as a hub by calculation of the betweenness centrality which is defined as the fraction of all shortest paths that go through a particular node [Dai et al., 2018]. Fig. 4c shows the distributions of betweenness centrality for the nodes in the S. cerevisiae networks, as well as the top 5 highest scoring nodes for this metric. Genes that have a high centrality in multiple networks are in bold print.

### 3.3 Cell type-specific networks in the mouse brain

In order to assess the similarity or difference between molecular networks in different cell types, networks were constructed from gene expression data from cells of six different neuronal and non-neuronal cell types in the mouse somatosensory cortex. Gene expression data was previously obtained using single-cell RNA-sequencing, and contain transcript counts of a total of 19972 genes in 1691 cells of the somatosensory cortex [Zeisel et al., 2015]. For computational efficiency, a random subset of 1000 genes was selected. Six cell-specific networks were constructed by using only cells from the particular type for network construction: astrocytes/ependymal (*N* = 143), endothelial/mural cells (*N* = 202), interneurons (*N* = 164), microglia (*N* = 84), oligodendrocytes (*N* = 699) and pyramidal cells (*N* = 399). The net-works were thresholded (*µ* + 3*σ*) after shuffling (see above) and sparsified. Fig. 5a shows the MI matrices of the cell type-specific networks after clustering using Principal Component Analysis (PCA). For each network, PCA was applied to the MI matrix and nodes were sorted by their highest-scoring component. This way, nodes with a similar connectivity profile are close to one another, revealing clustering in the cell-type specific molecular networks. The order of the nodes is different for each network, illustrating differences in local connectivity.

**Figure 5:**
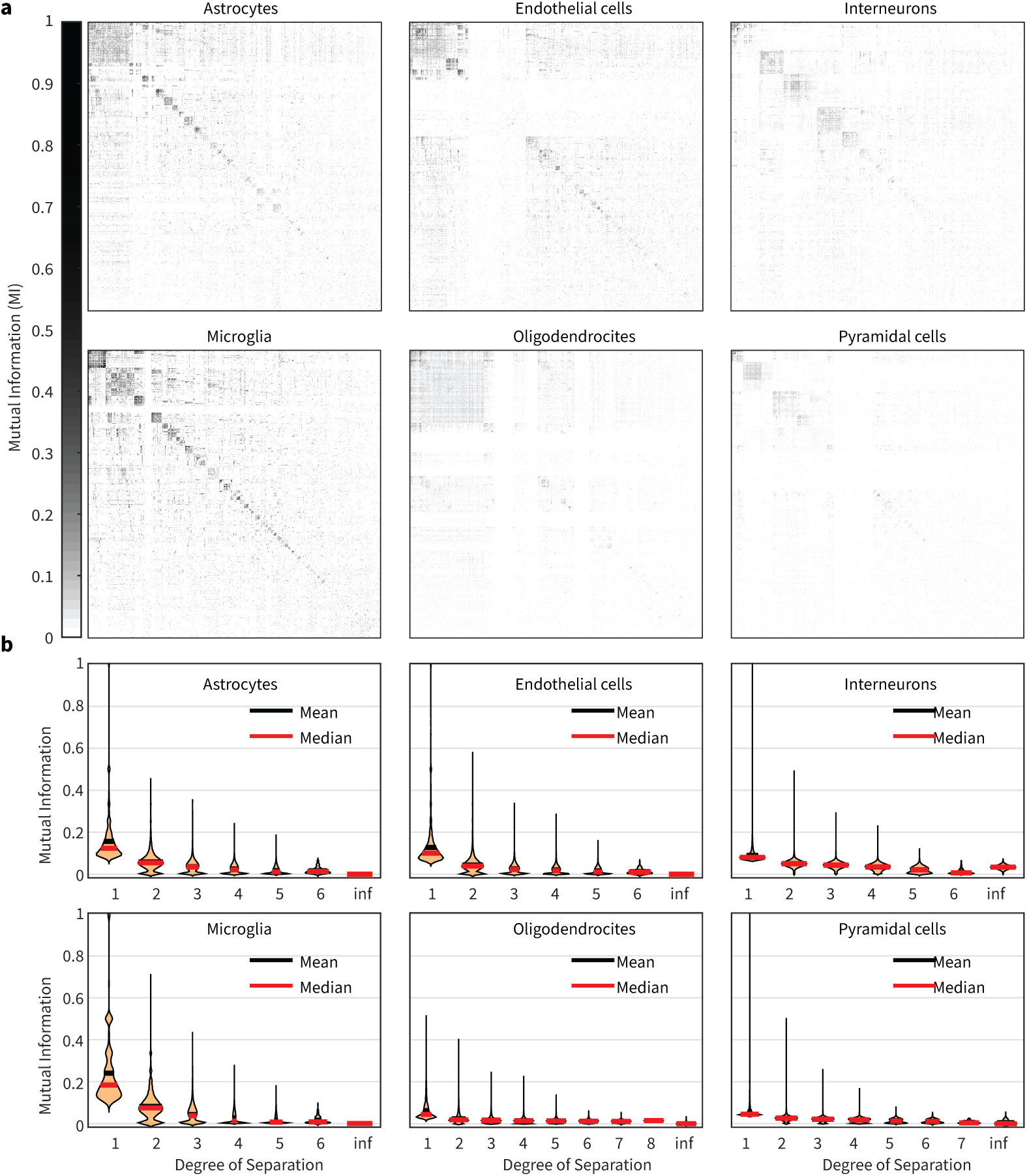
Cell type-specific gene co-expression networks in the mouse somatosensory cortex. **(a)** Mutual Information (MI) matrices showing pairwise correlations of genes in six different cell types in the somatosensory cortex. For every matrix PCA was used to sort the matrices, so that nodes with a similar connectivity profile are next to each other, illustrating differences in clustering between the networks. **(b)** MI distributions for increasing degree of separation (DOS) for each of the networks. A value of 1 means they are directly connected; a value of ‘inf’ means there is no possible path between them.

Similarly, to Fig. 2b for synthetic data, Fig. 5b shows the distributions of MI values as a function of DOS in cell type-specific networks. A DOS of ‘inf’ (infinite) occurs when there is no possible path between two nodes. The figures show that MI decreases as DOS increases, indicating that MI is indeed a measure of the ‘closeness’ of two nodes in the network, which decreases as noise is introduced with every added step along the path. Pathway analysis revealed for each node the betweenness centrality. Fig. 6 shows the distributions of these values for each network. Also displayed in Fig. 6 is, similar to Fig. 4c, the top 5 highest scoring nodes for this metric. Again, genes that are preserved across different networks are bold printed (note that this is not the case for any gene).

**Figure 6:**
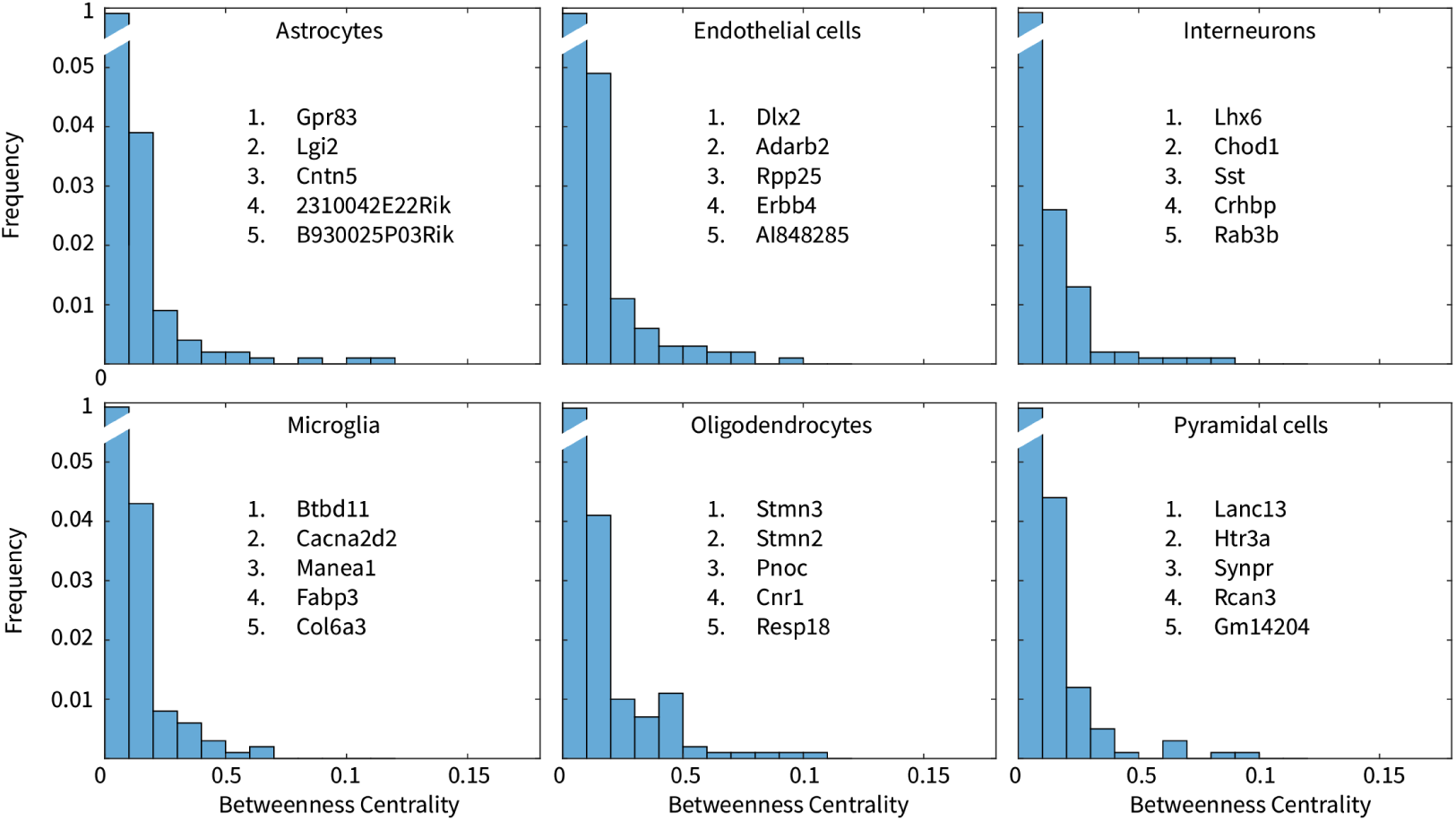
Cell type-specific gene co-expression networks in the mouse somatosensory cortex. Figures show the distributions of the nodes’ betweenness centrality in the six cell type-specific networks. Every plot lists the top five genes with the highest betweenness centrality. Genes that are preserved across different networks are bold printed.

### 3.4 Housekeeping Genes

Housekeeping Genes, also referred to as reference genes, are genes that are consistently ex-pressed in all cells and play essential roles in fundamental cellular functions required for maintaining cellular homeostasis. These genes maintain relatively stable expression levels across various tissue and experimental conditions, developmental stages, and environmental factors. In gene expression studies, these genes are commonly utilized as normalization factors to account for sample quality variations and RNA sequencing discrepancies.

To investigate the molecular networks of housekeeping genes in male and female mice, gene expression data from the somatosensory cortex layer 2*/*3 were utilized to construct networks. The gene expression data had been previously obtained using single-cell RNA sequencing and encompassed transcript counts for a total of 450 genes. The shared genes between males and females were identified, and gene-mutual information matrices were generated based on the gene expression data for each sex. Rows and columns containing solely 0 values were eliminated from both matrices. Fig. 7a displays the calculated mutual information matrices representing gene co-expression in the L2*/*3 somatosensory cortex of male and female mice. No sorting was performed on either of the matrices. The networks derived from these mutual information matrices were created by applying a threshold of (*µ* + *σ*) after shuffling (as described above) and subsequently sparsified. Fig. 7b presents the graph visualization obtained from the male mutual information matrix, exported as a network from NETSCOPE to Gephi following thresholding and sparsification. The node size corresponds to the gene’s expression count across all male samples. Fig. 7c illustrates the histogram of the betweenness centrality distribution for all genes, with each plot listing the top five genes exhibiting the highest betweenness centrality.

**Figure 7:**
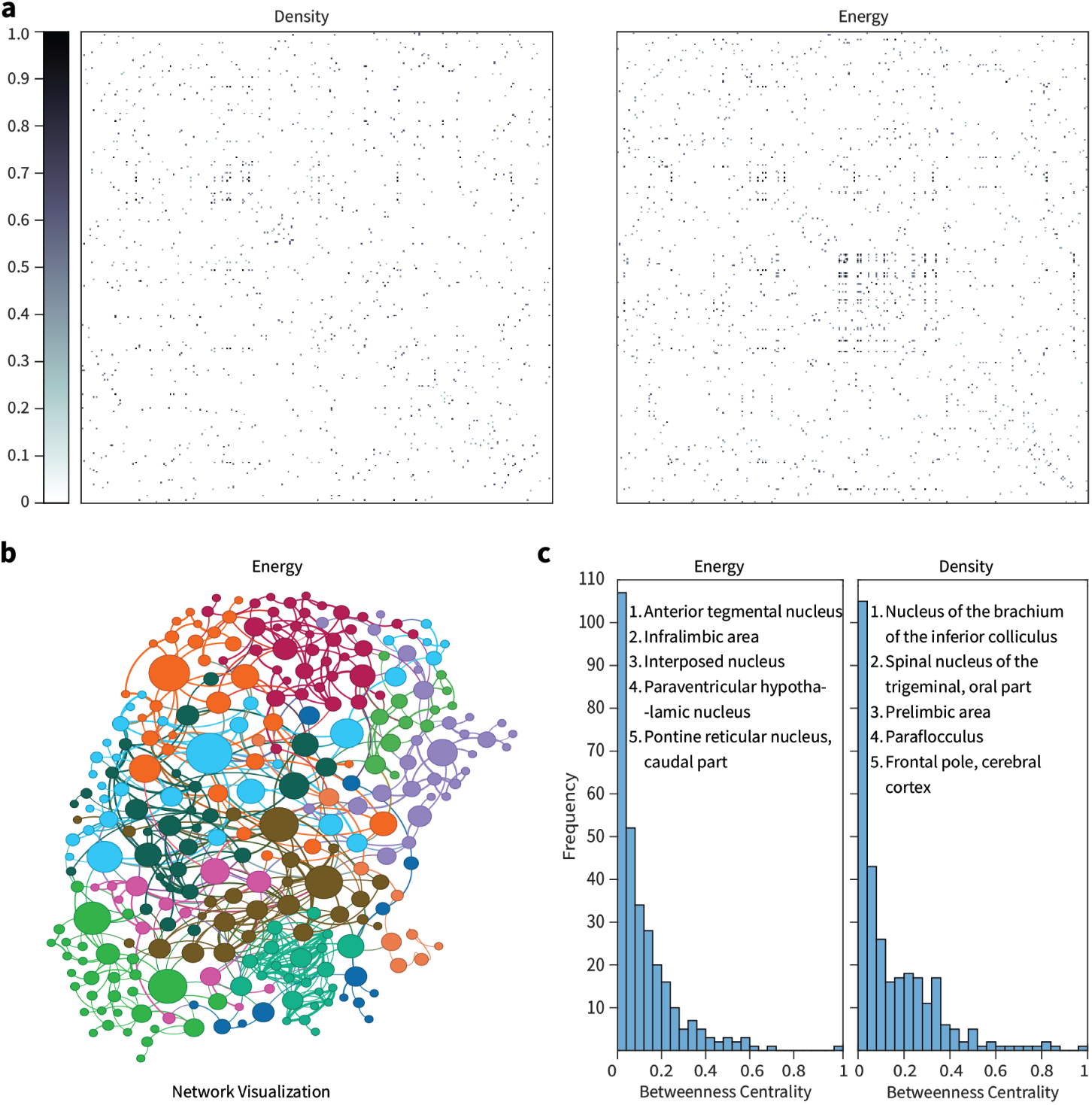
Housekeeping gene co-expression networks in the mouse somatosensory cortex. **(a)** Mutual Information (MI) matrices showing pairwise correlations of genes in Male and Fe-male mice, before network sparsification. **(b)** Visualization of the Male housekeeping gene correlation network generated from Gephi. The Mi matrix is thresholded using random shuffling, sparsified and then exported to Gephi. In Gephi, the **ForceAtlas2** network organization algorithm is applied. The edge between the two nodes represents the MI value between the genes. Independent nodes are removed. The the size of each node(4-40) are ranked using the betweenness centrality of the nodes. Colours depict the modularity classes that the nodes be-long to. **nooverlap** algorithm is applied so that no two nodes are overlapping. **(c)** Histogram of normalized Betweenness Centrality Male and Female housekeeping gene-Networks. The top five genes with the highest betweenness centrality are listed.

### 3.5 Cellular Networks

To investigate the mapping of transgenic lines in the somatosensory cortex, a comprehensive analysis was conducted using the Allen Brain Atlas. All transgenic line experiments targeting the somatosensory cortex were meticulously sorted, and the corresponding experiment numbers were exported to a CSV file. A total of 65 experiments were identified from the Allen Brain database and all relevant data associated with these experiments were exported to MATLAB. Subsequently, the data were merged into a single file and inputted into NETSCOPE for further processing.

In order to determine the transgenic line map, the data underwent processing in NETSCOPE, where mutual information (MI) was calculated for both Energy and Density measures across all 300 cells. Fig. 8a illustrates the resulting MI matrices, representing the co-expression patterns of cells in the somatosensory cortex of male and female mice. It is important to note that no sorting or any other manipulation was applied to these matrices. The networks were generated from these MI matrices by applying a threshold of (*µ* + *σ*) after shuffling, followed by sparsification.

**Figure 8:**
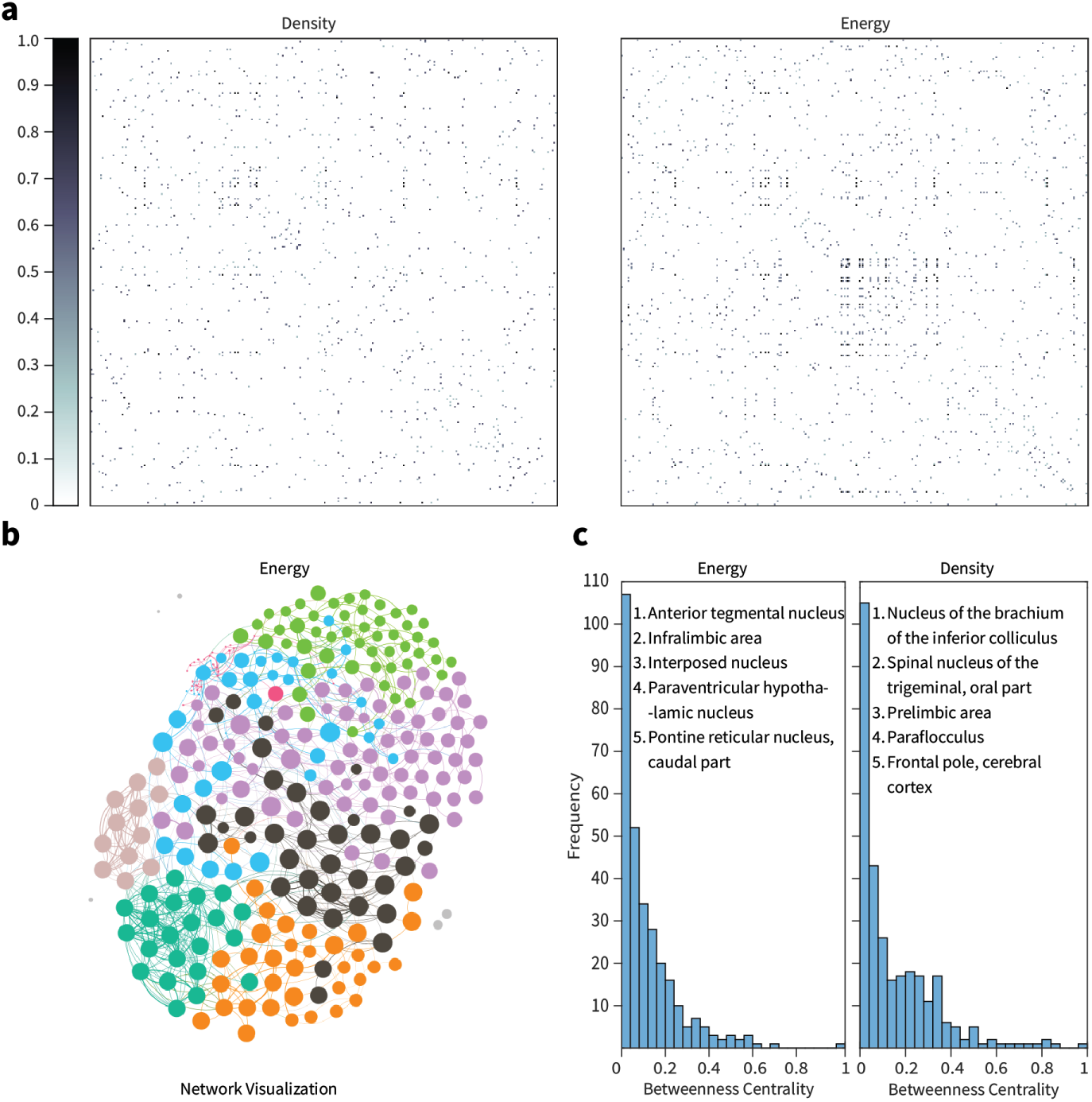
Cellular Networks in the mouse somatosensory cortex from Allen Brain Institute data. **(a)** Mutual Information (MI) matrices showing pairwise correlations of cells in the somatosensory cortex for Energy and Density of the injections, after thresholding (*µ* + *σ*) and network sparsification. **(b)** Visualization of the network for the energy generated from Gephi. The Mi matrix is exported to Gephi, **ForceAtlas2** network organization algorithm is applied. The edge between the two nodes represents the MI value between the genes. Independent nodes are removed. The size of each node(4-40) are ranked using the betweenness centrality of the nodes. The colours depict the modularity classes of the nodes. **nooverlap** algorithm is applied so that no two nodes are overlapping. **(c)** Histogram of normalized Betweenness Centrality for Energy and Density Networks. The top five cells with the highest betweenness centrality are listed.

Fig. 8b presents the graph visualization obtained from the Energy MI matrix, which was exported as a network from NETSCOPE to Gephi after undergoing thresholding and sparsification. The size of the nodes in the graph corresponds to the expression count of the gene across all male samples, providing a visual representation of the gene expression levels. Additionally, Fig. 8c displays a histogram showcasing the distribution of betweenness centrality for all genes. Each plot within the histogram highlights the top five genes with the highest betweenness centrality, indicating their significant role in information flow within the network.

### 3.6 Artificial network with timeseries

To demonstrate that NETSCOPE can reconstruct networks from continuous-valued data, we generated a synthetic EEG dataset consisting of 32 channels. A ground-truth network was constructed to reflect biologically plausible properties, including sparsity and small-world organization. Using simulated continuous time series, NETSCOPE was applied to infer the underlying connectivity structure. The results show that NETSCOPE successfully reconstructs the ground-truth network (See Fig. 9 a and b, the method remains robust under increasing noise levels: with 10% additive noise, NETSCOPE achieves a TPR exceeding 80% (See Fig. 9 c). Even at 25% additive noise, the TPR remains around 60%, while maintaining a false positive rate FPR below 20%. Although reconstruction performance decreases with higher noise levels, NETSCOPE continues to recover meaningful network structure, albeit with an increased false positive rate under stronger noise contamination. Notably the TPR connections maintain higher MI values.

**Figure 9:**
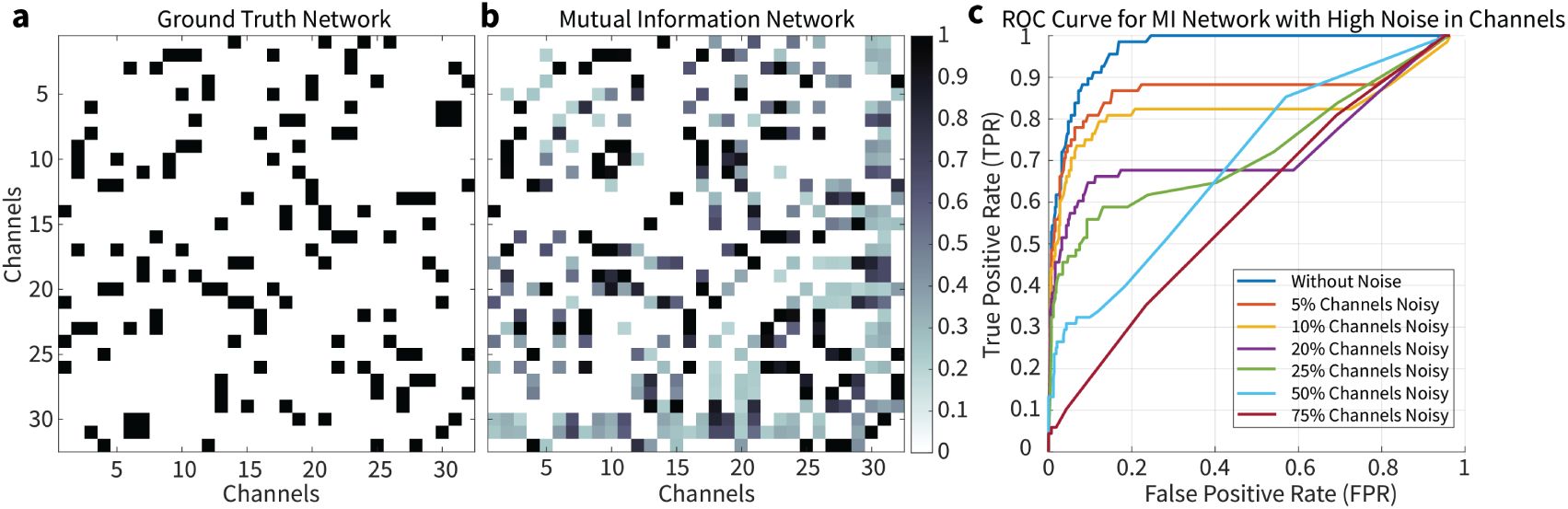
**(a)** Ground Truth Network of the artificial network. This connection matrix is used for creating the artificial eeg signals. The connection matrices shows the small world property. **(b)** The reconstructed connection matrix using NETSCOPE after thresholding. **(c)** ROC curve, when a high level of noise is introduced on the EEG signals. the different colors shows home many of the EEG signals were given additional high noise.

### 3.7 EEG Networks

The application of the NETSCOPE framework to EEG recordings revealed distinct and temporally evolving functional connectivity patterns across the four defined conditions. The mutual information (MI)-based networks captured non-linear dependencies between EEG channels, enabling a detailed characterization of functional interactions.

During the *auditory baseline* period, the network exhibited relatively low overall connectivity, with sparse interactions primarily distributed across frontal and central regions (See Fig. 10a). This pattern is consistent with a resting-state configuration with limited coordinated activity prior to stimulus presentation.

**Figure 10:**
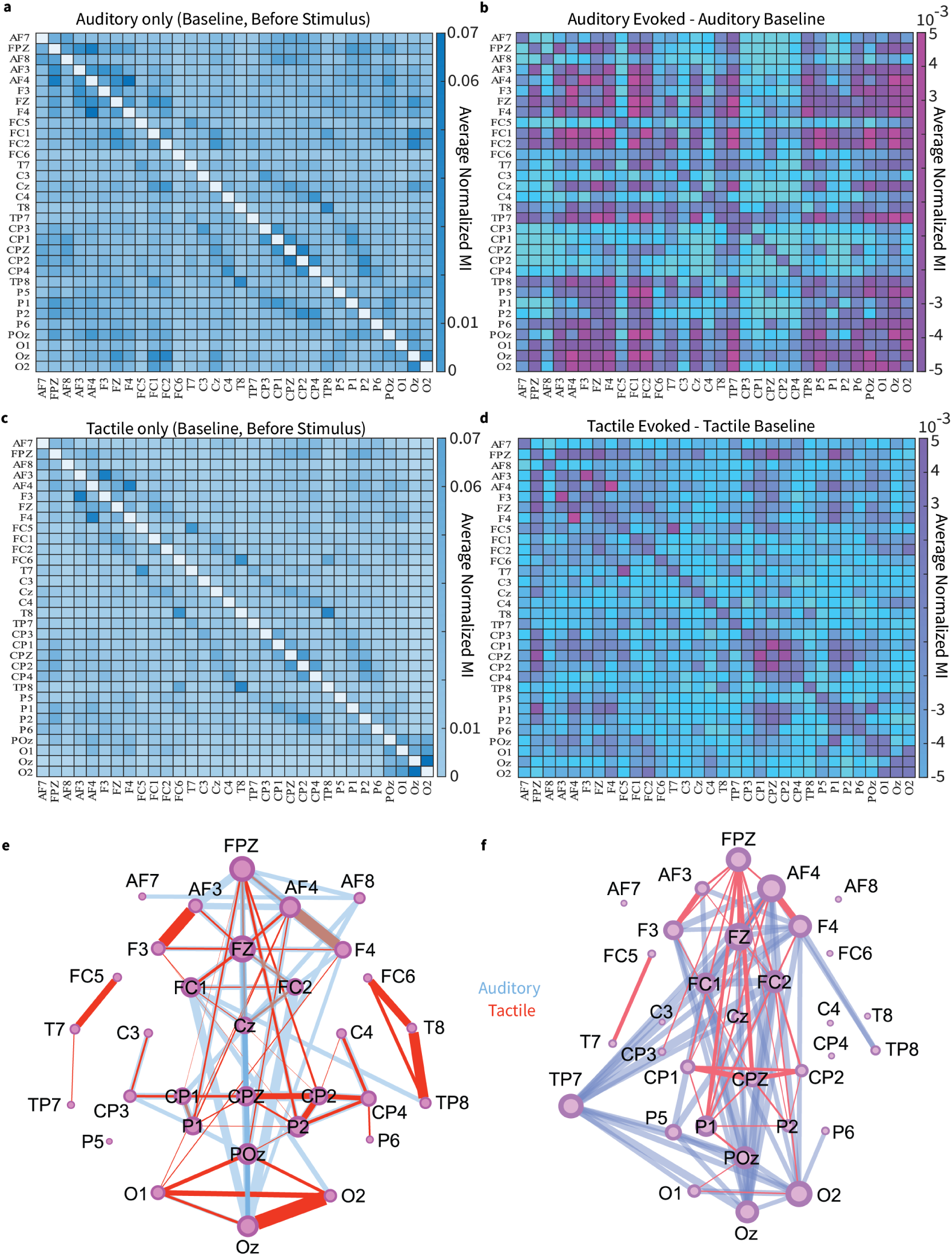
With the NETSCOPE toolbox, MI matrices between the 32 channels from the time frame of interest can be calculated. In the exploratory analysis carried out in this example, four time frames are chosen for the analysis: (1)250 ms before the auditory stimulus onset, (2)250 ms before the tactile stimulus onset, (3)250 ms after the auditory stimulus onset and (4)400 ms after the tactile stimulus onset. The above mentioned time frame were conceptualized as (1)the auditory baseline (2)the tactile baseline22(3)the auditory evoked potential (4)the tactile evoked potential. Four MI matrices were calculated using the NETSCOPE toolbox from those time frames of interest. The top 10% edges are then filtered out and carried over to Gephi for network visualization.

In the *auditory evoked* condition, a clear increase in connectivity strength was observed, particularly involving frontocentral and temporal electrodes. The network showed a more organized structure, with the emergence of hub-like nodes in frontal regions(See Fig. 10b), suggesting their involvement in early sensory processing and attentional modulation.

The *tactile baseline* condition displayed a network structure comparable to the auditory baseline, but with slightly enhanced connectivity in central and parietal regions(See Fig. 10c). This may reflect anticipatory processes related to the expected tactile stimulus.

The *tactile evoked* network exhibited the most pronounced reorganization. A substantial increase in connectivity was observed, especially among central (C_Z_, C_3_, C_4_) and parietal electrodes, consistent with somatosensory processing. The resulting network was more integrated and denser within the retained top 10% of edges, indicating stronger synchronization across distributed cortical regions(See Fig. 10d).

Importantly, the MI-based approach produced sparse yet highly informative networks, retaining only the strongest 10% of connections while preserving meaningful functional structure. This sparsification enhances interpretability by emphasizing dominant interactions and reducing noise-driven connections(See Fig. 10e and f).

Comparative analysis across conditions revealed a consistent transition from baseline to stimulus-evoked states, characterized by increased global connectivity, the emergence of condition-specific hubs, and enhanced integration within task-relevant cortical regions.

Overall, these results demonstrate that NETSCOPE effectively captures dynamic, stimulus-dependent reorganization of brain networks and highlights the utility of mutual information for detecting non-linear and transient functional interactions in EEG data.

### 3.8 Functional network

The comparison between mutual information (MI) and Pearson correlation-based functional connectivity (FC) revealed both overlapping and distinct patterns of brain connectivity, highlighting the complementary nature of linear and nonlinear measures.

To facilitate interpretation, the edge overlap matrix encodes connectivity differences as follows: a value of 1 indicates edges present only in the FC network, a value of 2 indicates edges present only in the MI network, and a value of 3 denotes edges shared between both methods. This representation enables direct visualization of common and method-specific connections.

The edge overlap matrix (Fig. 11a) demonstrated that a substantial subset of connections was shared between FC and MI, indicating that strong linear correlations often coincide with nonlinear dependencies. However, a significant proportion of edges were uniquely identified by MI, suggesting that nonlinear interactions capture additional functional relationships not detectable through Pearson correlation alone.

**Figure 11:**
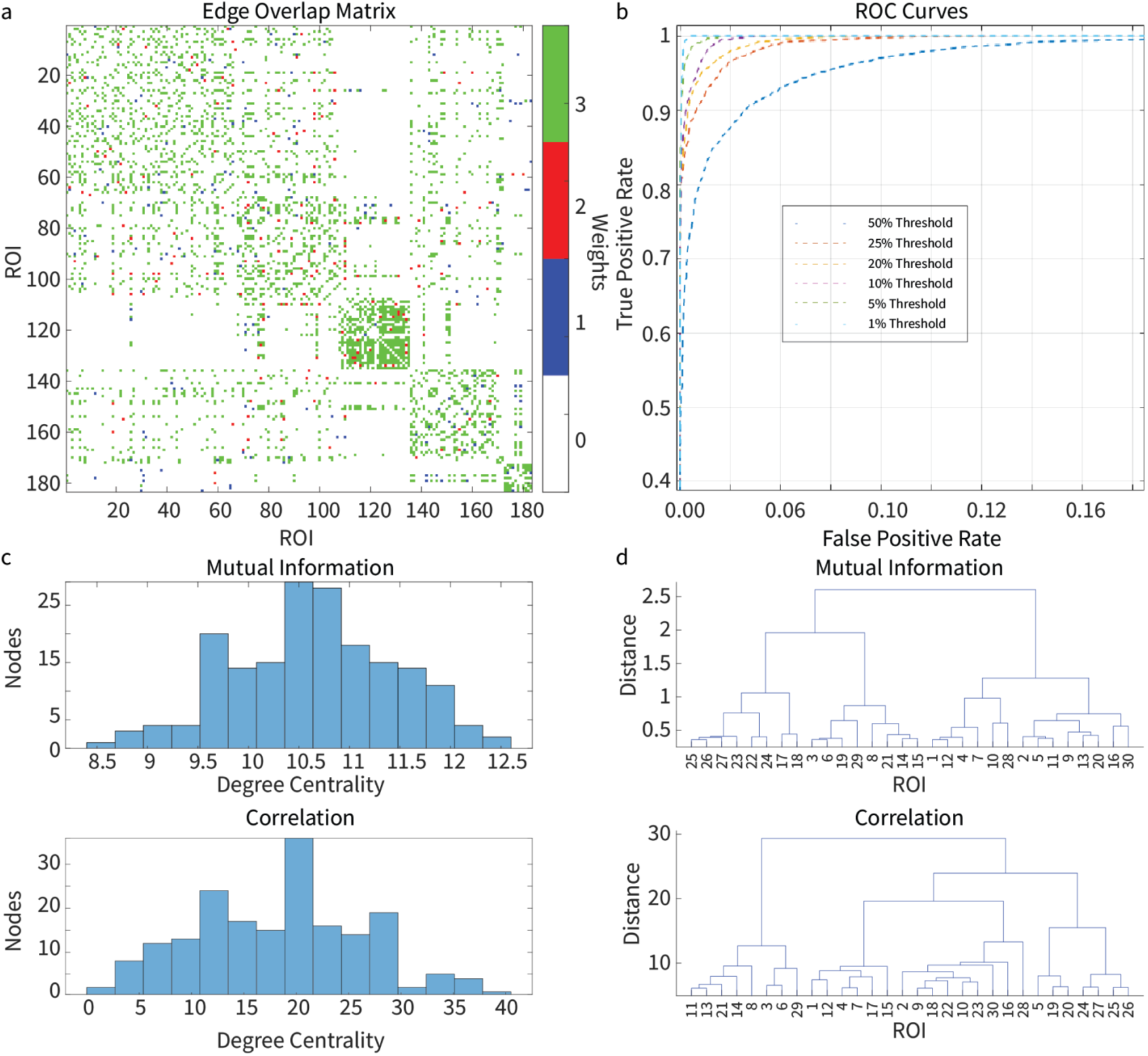
**(a)** Edge Overlap Matrix: A comparison of functional connectivity (FC) and mutual information (MI) connectivity matrices using the top 10% of connections from the Pearson’s correlation-based FC matrix as a threshold for significant edges. Binary connectivity matrices were generated for both FC and MI, assigning values of 1 to FC-specific edges, 2 to MI-specific edges, and 3 to edges shared by both methods. The resulting matrix visualizes overlapping and distinct connectivity features, emphasizing the complementary roles of linear (Pearson’s correlation) and nonlinear (mutual information) measures in capturing brain network interactions. **(b)** ROC Curve: Evaluation of NETSCOPE’s performance across multiple Pearson’s correlation thresholds (1%, 5%, 10%, 20%, 25%, and 50%), demonstrating its ability to detect significant edges and assess reproducibility in functional connectivity. **(c)** The histogram of weighted and unweighted degree centrality. **(d)** Dendogram of the networks created by correlation and MI methods

Notably, MI-specific connections were more broadly distributed across cortical and subcortical regions, reflecting the ability of MI to detect complex and potentially higher-order dependencies. In contrast, FC-specific edges were more localized and predominantly reflected strong linear synchronization patterns.

The ROC analysis (Fig. 11b) demonstrated that the agreement between MI and FC varies as a function of the threshold applied to the FC matrix. At lower thresholds, a larger number of connections were included, resulting in increased overlap but reduced specificity. As the threshold increased, the overlap decreased, indicating that MI retains connections that are not captured even among the strongest linear correlations.

Furthermore, network-level analysis revealed that MI-based networks exhibited distinct topological properties compared to FC networks. Specifically, MI networks showed increased integration and a more distributed connectivity profile, whereas FC networks were characterized by stronger clustering around highly correlated regions. The inclusion of edge weights further emphasized these differences, with weighted MI networks preserving subtle interaction strengths that are not captured in binarized representations (See Fig. 11c, d).

## 4 Discussion

Here we developed NETSCOPE, a multiplatform (Python/MATLAB/Octave) toolbox that can be used for the network discovery and reconstruction of networks. NETSCOPE takes advantage of mutual information (MI) and Variation of Information (VI) to uncover correlations from molecular to functional networks.

Traditional approaches to analyzing brain networks have relied heavily on correlation-based measures, such as Pearson’s correlation coefficient, to quantify the relationships between different network components. Although these measures are useful for identifying pairwise relationships, they have several limitations. For instance, correlation-based measures are sensitive to linear relationships, making them less effective at capturing nonlinear interactions that are characteristic of brain networks. Additionally, correlation-based measures are symmetric, making it difficult to infer directional relationships between network components. Furthermore, correlation-based measures are often susceptible to false positives, particularly in high-dimensional datasets.

In contrast, information-theoretic measures, such as mutual information and variation of information, offer a more nuanced understanding of network interactions. These measures are better suited for handling the nonlinear, high-dimensional relationships that exist within and between brain networks. Information theoretic measures can capture directional relationships to infer causal interactions between network components. Moreover, information-theoretic measures are more robust to false positives, providing a more accurate representation of network organization. However, information-theoretic measures are not without their limitations. One major concern is bias, which can arise from the estimation of probability distributions from finite samples. This bias can lead to inaccurate estimates of mutual information and variation of information, particularly when dealing with high-dimensional datasets (see Fig.3). Additionally, information-theoretic measures can be sensitive to sampling variability, which can result in inconsistent estimates of network interactions (see Fig. 3).

The unique properties of MI and VI enabled a rapid and robust data processing pipeline, making NETSCOPE applicable for studying well-sampled biological networks, e.g. for network discovery (see Supplementary Fig. 1 and Supplementary Fig. 2, network reconstruction (Supplementary Fig. 6) and differential network regulation (see Supplementary Fig. 3). To validate the effectiveness of the toolbox, we compared NETSCOPE performance against ground truth, using synthetic data with known network topology, ensuring its sensitivity and selectivity. Furthermore, we reconstructed transcriptional networks in different yeast species [Ziemann et al., 2019] and compared the results against established genetic networks in yeast.

NETSCOPE works in a plug-and-play manner without the need for additional toolbox installation or complicated setup; the code is available on GitHub (links comes here) and the expression data can be imported into Colab, Python, Octave or MATLAB from e.g. a comma-separated value (CSV) file. The toolbox offers the flexibility to fine-tune any parameters in the data processing pipeline but also works out of the box using default parameters - the latter of which was the case for the analyses described in this article. Network data can be exported to network software tools for visualization and subsequent analysis (see Supplementary Figs. 5 and 6).

While MI is computationally heavy, parallelization can speed up the process. For the parallel computation of the MI matrix, one can use the MATLAB Parallel Processing Toolbox in combination with the parallel processing functions included in NETSCOPE. Due to the universal nature of MI, NETSCOPE allows for the cross-comparison of transcription data between cell types, individuals, populations, or species. Rather than comparing individual genes, one can now compare entire transcriptomes by their internal structure, looking for patterns of connectivity that are preserved [Breschi et al., 2017]. These patterns can, for example, shine a light on basic molecular mechanisms at the root of the evolutionary tree, which would be imperceptible through single gene analysis.

Going beyond cross-species transcriptomics, NETSCOPE can be used to integrate multi-omics data covering multiple layers of the cellular process - e.g. genomics, epigenomics, proteomics and metabolomics - to reconstruct network representations of the molecular landscape. Understanding the topology of these networks makes it possible to bridge the gap between genotype and phenotype and predict the downstream effects of genetic defects [Ma et al., 2020]. It will allow us to link low-level physiological mechanisms to higher-level phenomena such as disease symptoms and behavioral traits, and provide insight into the mechanisms behind cell fate and how to control it, allowing for cellular reprogramming.

---

**Supplementary Figure 1:**
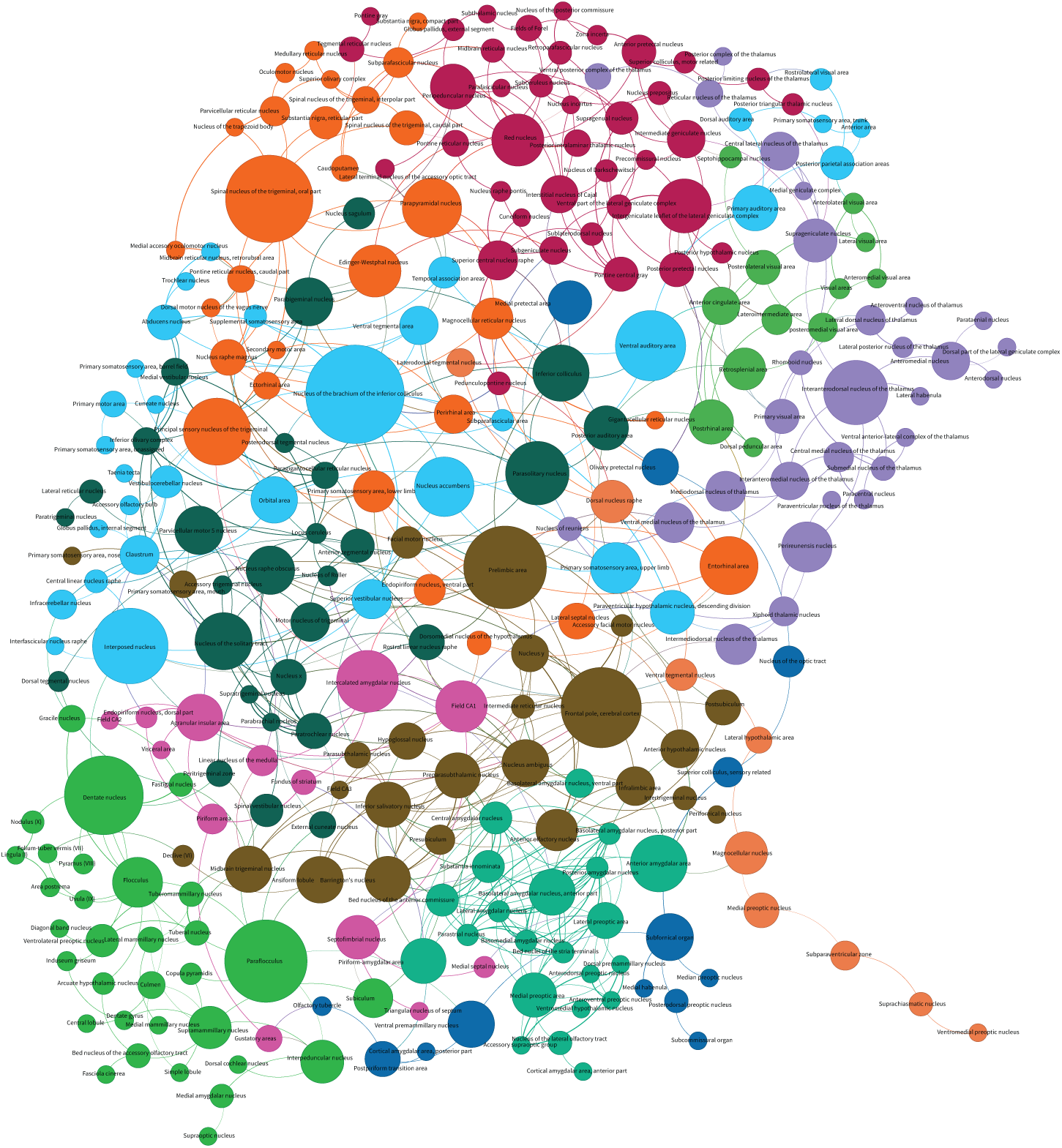
Visualization of Density-network of the cellular network created from Allen Brain Atlas data. **a** Visualization of the Male housekeeping gene correlation net-work with all the gene names generated from Gephi. All positive values from the differential Mi matrix are sorted to get the gene co-expression matrix enriched in the male samples. The matrix is thresholded using random shuffling and sparsified before exporting to Gephi. In Gephi, the **ForceAtlas2** network organization algorithm is applied. The edge between the two nodes represents the MI value between the genes. Independent nodes are removed. The colour and the size of each node(4-40) are ranked using the betweenness centrality of the nodes. **nooverlap** algorithm is applied so that no two nodes are overlapping

**Supplementary Figure 2:**
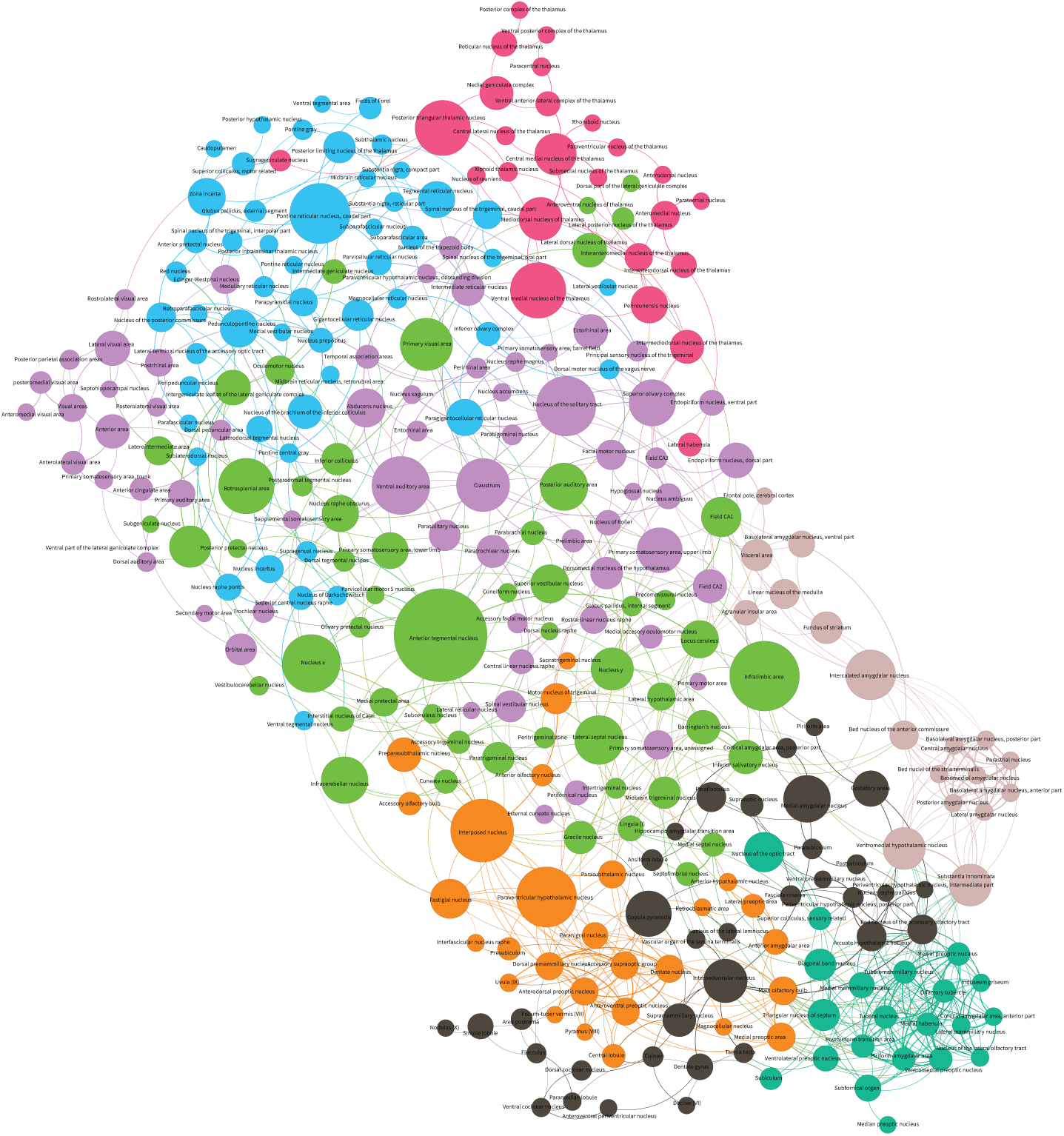
Visualization of Energy-network of the cellular network created from Allen Brain Atlas data. **a** Visualization of the Male housekeeping gene correlation network with all the gene names generated from Gephi. All positive values from the differential Mi matrix are sorted to get the gene co-expression matrix enriched in the male samples. The matrix is thresholded using random shuffling and sparsified before exporting to Gephi. In Gephi, the **ForceAtlas2** network organization algorithm is applied. The edge between the two nodes represents the MI value between the genes. Independent nodes are removed. The colour and the size of each node(4-40) are ranked using the betweenness centrality of the nodes. **nooverlap** algorithm is applied so that no two nodes are overlapping

**Supplementary Figure 3:**
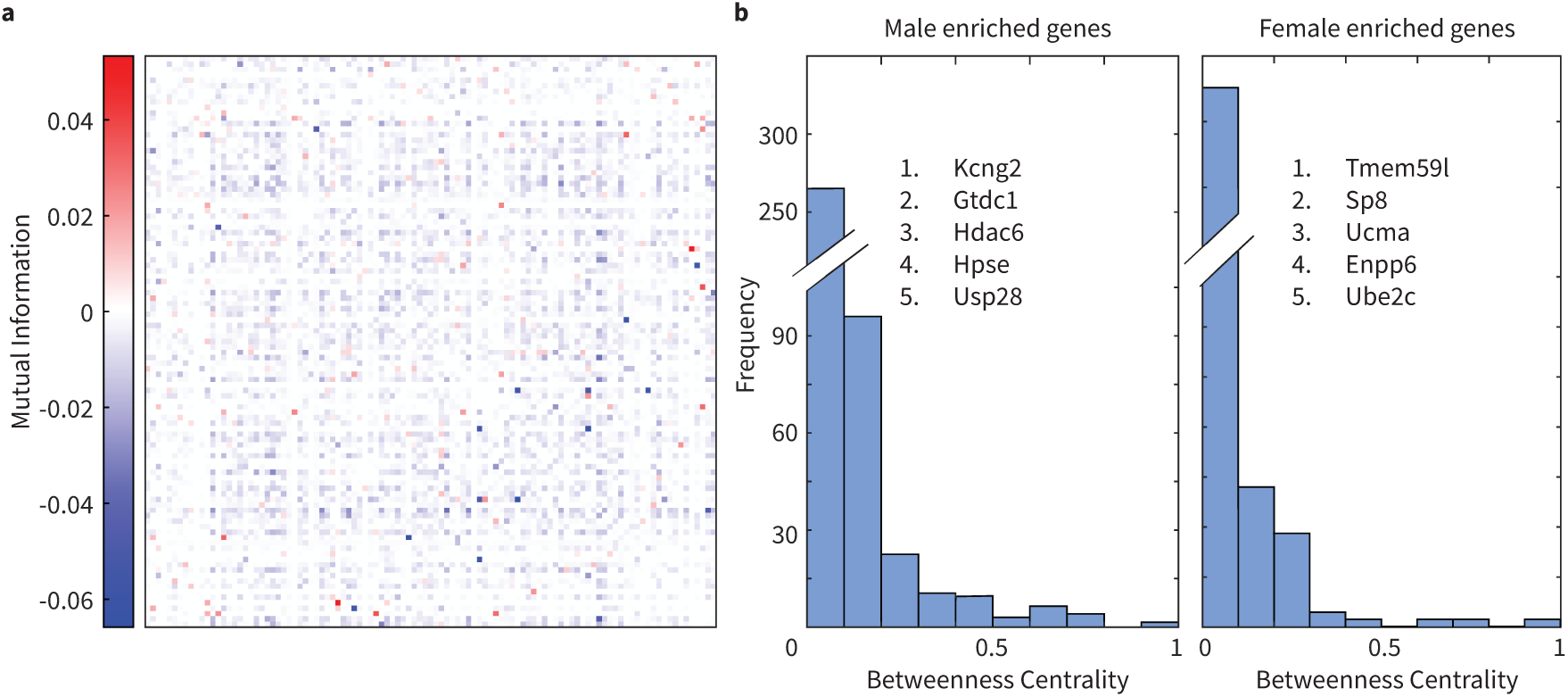
Differential network for Male and Female. **(a)** Mutual Information (MI) matrix showing the difference of the housekeeping gene MI matrices for Male and Female. **(b)** Histogram of normalized Betweenness Centrality of the network for male enriched genes(i.e. positive values in differential MI matrix) and female enriched genes(i.e. negative values in the differential MI matrix). The top five genes with the highest betweenness centrality are listed.

**Supplementary Figure 4:**
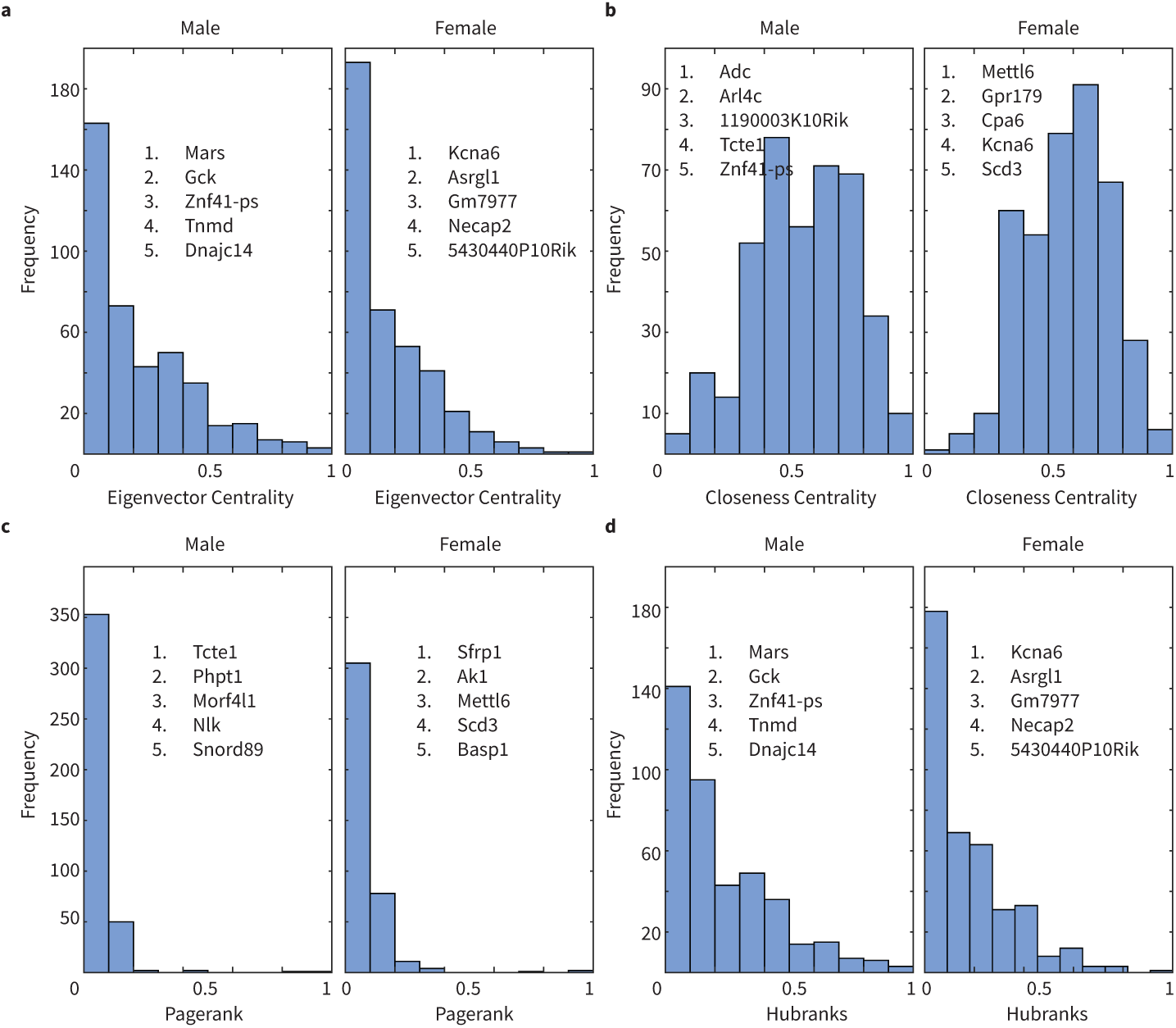
Network measures of networks generated by NETSCOPE after thresholding and Sparsifying the MI matrix of Housekeeping genes in the mouse somatosensory cortex. **(a)** Histogram of the eigenvector centrality for Male and Female house-keeping gene MI networks.**(b)** Histogram of the closeness centrality for Male and Female housekeeping gene MI networks. **(c)** Histogram of the normalized pagerank for Male and Female housekeeping gene MI networks. **(d)** Histogram of the normalized hubranks for Male and Female housekeeping gene MI networks. The top 5 genes with the highest values are listed in each figure.

**Supplementary Figure 5:**
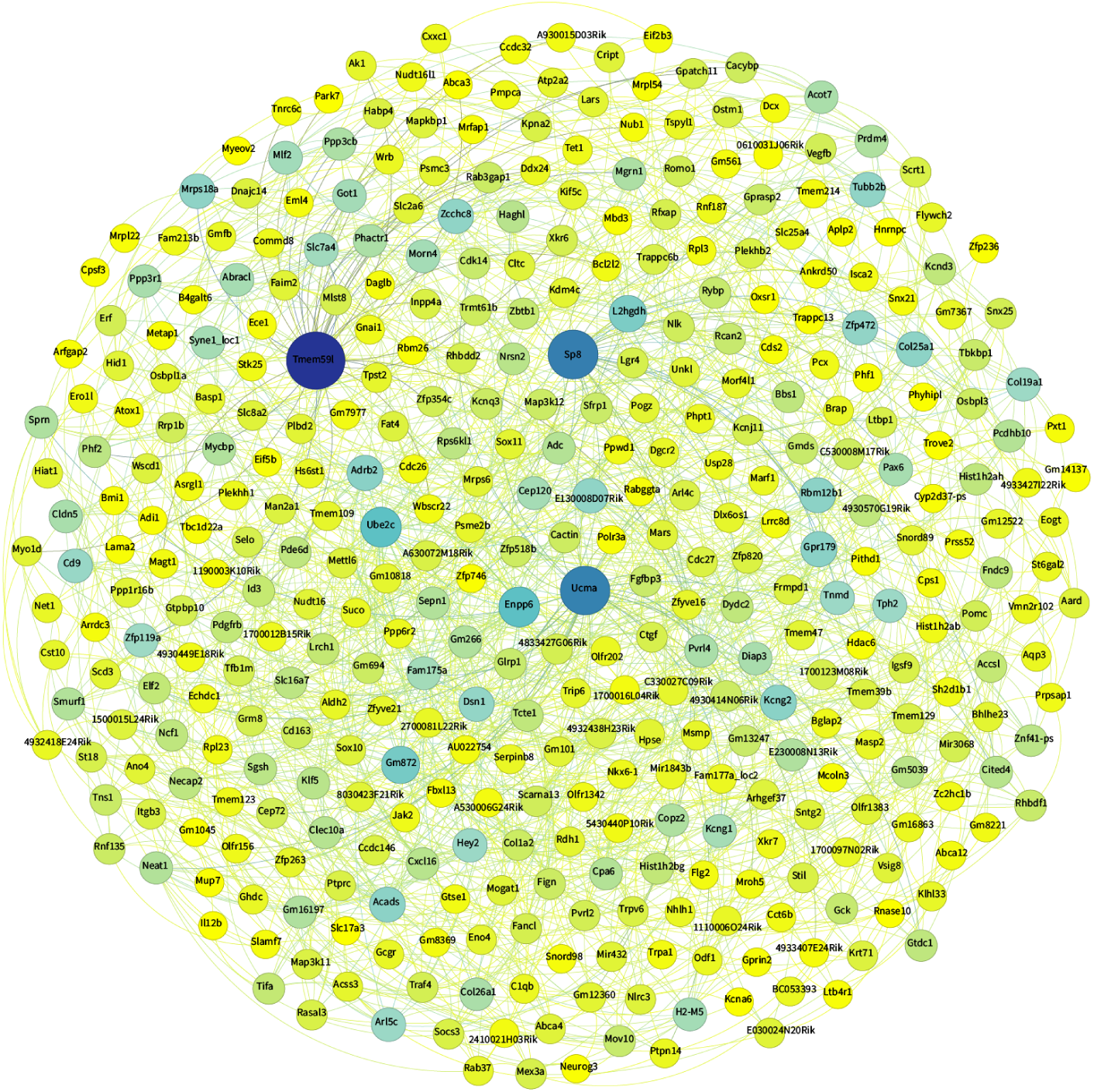
Housekeeping gene co-expression networks in the mouse somatosensory cortex. **(a)** Visualization of the Male housekeeping gene correlation network with all the gene names generated from Gephi. All positive values from the differential Mi matrix are sorted to get the gene co-expression matrix enriched in the male samples. The matrix is thresholded using random shuffling and sparsified before exporting to Gephi. In Gephi, the **ForceAtlas2** network organization algorithm is applied. The edge between the two nodes rep-resents the MI value between the genes. Independent nodes are removed. The colour and the size of each node(4-40) are ranked using the betweenness centrality of the nodes. **nooverlap** algorithm is applied so that no two nodes are overlapping

**Supplementary Figure 6:**
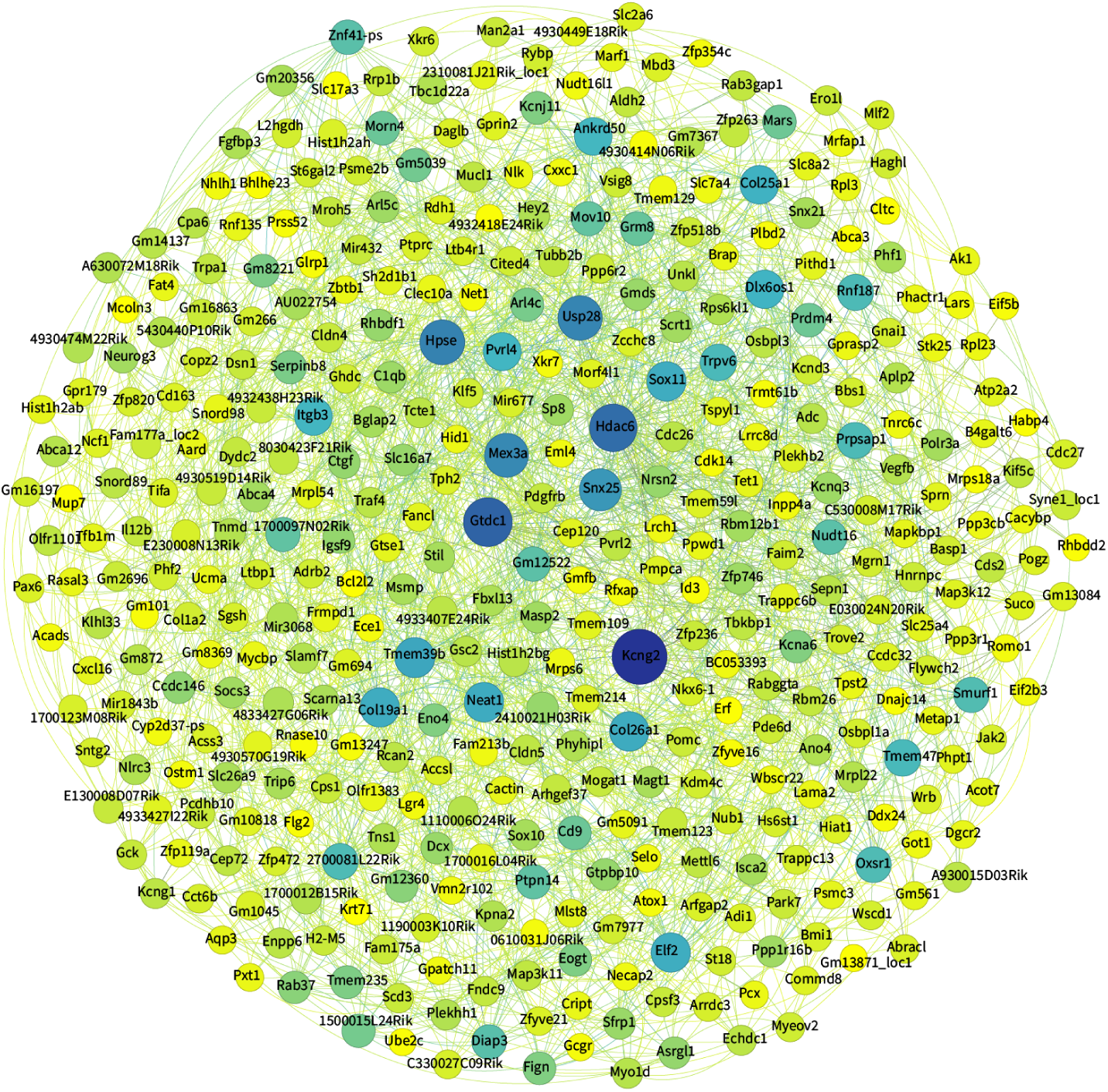
Housekeeping gene co-expression networks in the mouse somatosensory cortex. **(a)** Visualization of the Male housekeeping gene correlation network with all the gene names generated from Gephi. All positive values from the differential Mi matrix are sorted to get the gene co-expression matrix enriched in the male samples. The matrix is thresholded using random shuffling and sparsified before exporting to Gephi. In Gephi, the **ForceAtlas2** network organization algorithm is applied. The edge between the two nodes rep-resents the MI value between the genes. Independent nodes are removed. The colour and the size of each node(4-40) are ranked using the betweenness centrality of the nodes. **nooverlap** algorithm is applied so that no two nodes are overlapping

